# Ezh2 as an epigenetic checkpoint regulator during monocyte differentiation: a potential target to improve cardiac repair after myocardial infarction

**DOI:** 10.1101/2021.02.17.428828

**Authors:** Rondeaux Julie, Groussard Déborah, Renet Sylvanie, Tardif Virginie, Dumesnil Anaïs, Chu Alphonse, Henry Jean-Paul, Badji Zina, Vézier Claire, Béziau Delphine, Guerrot Dominique, Brand Marjorie, Richard Vincent, Durand Eric, Brakenhielm Ebba, Fraineau Sylvain

## Abstract

Epigenetic regulation of histone H3K27 methylation has recently emerged as a key step during M2-like macrophage polarization, essential for cardiac repair after Myocardial Infarction (MI). We demonstrate for the first-time that EZH2, responsible for H3K27 methylation, has an ectopic cytoplasmic localization during monocyte differentiation in M2 macrophages. Moreover, we show that pharmacological EZH2 inhibition, with GSK-343, enhances bivalent genes, expression to promote human monocyte repair functions. GSK-343 treatment accelerated cardiac inflammatory resolution preventing infarct expansion and subsequent cardiac dysfunction after MI *in vivo*. In conclusion, our study reveals that epigenetic modulation of cardiac-infiltrating immune cells may hold promise to limit adverse cardiac remodeling after MI.

## Main

Myocardial infarction (MI) triggers an endogenous wound healing response involving an early innate immune response starring myeloid cells. Macrophages are the predominant myeloid cell type arising from bone marrow-mobilized circulating monocytes implicated in post-MI cardiac repair^1^. Rapidly after MI, circulating classical, pro-inflammatory Ly6C^hi^ monocytes are recruited to the heart and accumulate at the ischemic area where they differentiate into pro-inflammatory M1 macrophages and give rise to the early inflammatory phase within the first 3 days post-MI in mice. Over the course of cardiac repair, during the first week after MI, the inflammatory phase gives way to the repair phase, characterized by a switch of pro-inflammatory M1 macrophages toward an alternative, immunomodulatory Ly6C^lo^ M2 macrophage phenotype^2^. Although cardiac-recruited Ly6C^hi^ monocytes initially differentiate towards M1 macrophages, they gradually differentiate also into reparative M2 macrophages^2^. Importantly, this macrophage phenotype transition promotes angiogenesis, lymphangiogenesis, extracellular matrix synthesis and deposition necessary to create a mature fibrotic scar. The proper sequence of inflammatory and reparative phases is critical to prevent cardiac remodeling and dysfunction to limit development of heart failure. Experimental studies have shown that either reduced cardiac monocyte infiltration^3^ and absence of a proper pro-inflammatory phase^4^ or sustained duration of the acute inflammatory phase due to M1 macrophage accumulation^5^, but also reduced monocyte to M2 macrophage differentiation during the reparative phase^6^, ^7^ all result in defective post-MI cardiac healing. Indeed, rapid resolution of the inflammatory phase, giving way to an early and strong reparative phase, correlates with reduction of infarct scar expansion and better preservation of cardiac function.

In this context, macrophage phenotype switching, from M1 to M2-like, plays a key role in cardiac inflammation and repair post-MI. Accumulating evidence points to a specific epigenetic histone modification, histone H3 lysine 27 trimethylation (H3K27me3), as an important mechanism to regulate macrophage activation and polarization^8-10^. The repressive H3K27me3 mark is generated by the Polycomb repressive complex 2 (PRC2), containing the epigenetic histone methyltransferase enzyme Enhancer of zeste homolog 2 (Ezh2). Conversely, H3K27 methylation is actively antagonized by two histone demethylases: Jumonji domain-containing protein D3 (Jmjd3) and Ubiquitously transcribed TPR on X (Utx). This fine-tuned balance between activators and repressors of epigenetic modifications on H3K27 impacts the expression of many genes involved in diverse biological processes including cell-fate commitment during cellular differentiation. Jmjd3 has been described as an essential regulator of M2 macrophage polarization through upregulation of *Arg1 Chi3l3*, *Retnla* and *CCL17* linked to *Irf4* up-regulation during helminth infection^11^ as well as following IL4 stimulation in both human monocytes^12^ and murine macrophages^13^. Although Jmjd3 implication during M2 macrophage polarization is well established, the role of its antagonistic partner, Ezh2, has not yet been well established in macrophages. In contrast, Ezh2 has been suggested to play important roles in regulating inflammatory cells functions in B cells^14^, ^15^, T cells^16^, neutrophils and dendritic cells^17^. However, recent reports suggest that Ezh2 may influence macrophage activation by suppressing the expression of anti-inflammatory genes^18^ and reducing cytokine secretion leading to a reduction of macrophage-dependent disease development^19^, ^20^.

Altogether, these findings suggest that H3K27 methylation promotes a M1-like pro-inflammatory state, while H3K27 demethylation promotes a M2-like immunomodulatory phenotype in macrophages.

We hypothesized that Ezh2, by modulating H3K27 methylation, may directly regulate monocyte to M2 macrophage differentiation and thus represents an attractive therapeutic target to design new pharmacological epigenetic inhibitors to promote cardiac repair post-MI and prevent the occurrence of heart failure. Here we demonstrate that Ezh2 serves as an epigenetic checkpoint regulator during monocyte to macrophage M2 differentiation, by translocating from the nucleus to the cytoplasm, leading to derepressed expression of bivalent genes essential for M2-like macrophage differentiation and function. Moreover, we promisingly discovered that pharmacological Ezh2 inhibition, with GSK-343, reduced the H3K27me3 mark specifically on the promotor of bivalent genes (such as *DLL1*, *VEGFA*, *IRF4*) resulting in accelerated resolution of the inflammatory phase leading to reduced infarct scar expansion and improved cardiac function after MI in mice.

## Results

### Cytoplasmic translocation of Ezh2 in cardiac M2-like macrophages post-MI

We first investigated Ezh2 localization in cardiac-infiltrating myeloid cells in mice during post-MI cardiac repair. As expected, we observed a nuclear localization of Ezh2 in both CD11b^+^/CD68^-^ and Cd11b^+^/Cd68^+^ cardiac monocytes (Fig. 1a) and Cd68^+^/iNos^+^ cardiac M1 differentiated macrophages (Fig. 1b). Unexpectedly, Ezh2 translocated to the cytoplasm in cardiac differentiated post-MI (Fig. 1c) as well as in cardiac-resident (Fig. 1d) Cd68^+^/Cd206^+^ M2 macrophages in healthy mice. These observations suggest that cytoplasmic translocation of Ezh2 might promote M2-like macrophage differentiation by initiating a switch from pro-inflammatory M1 toward immunoregulatory M2 phenotype and subsequent transition from pro-inflammatory toward reparative phase after MI.

**Figure 1:**
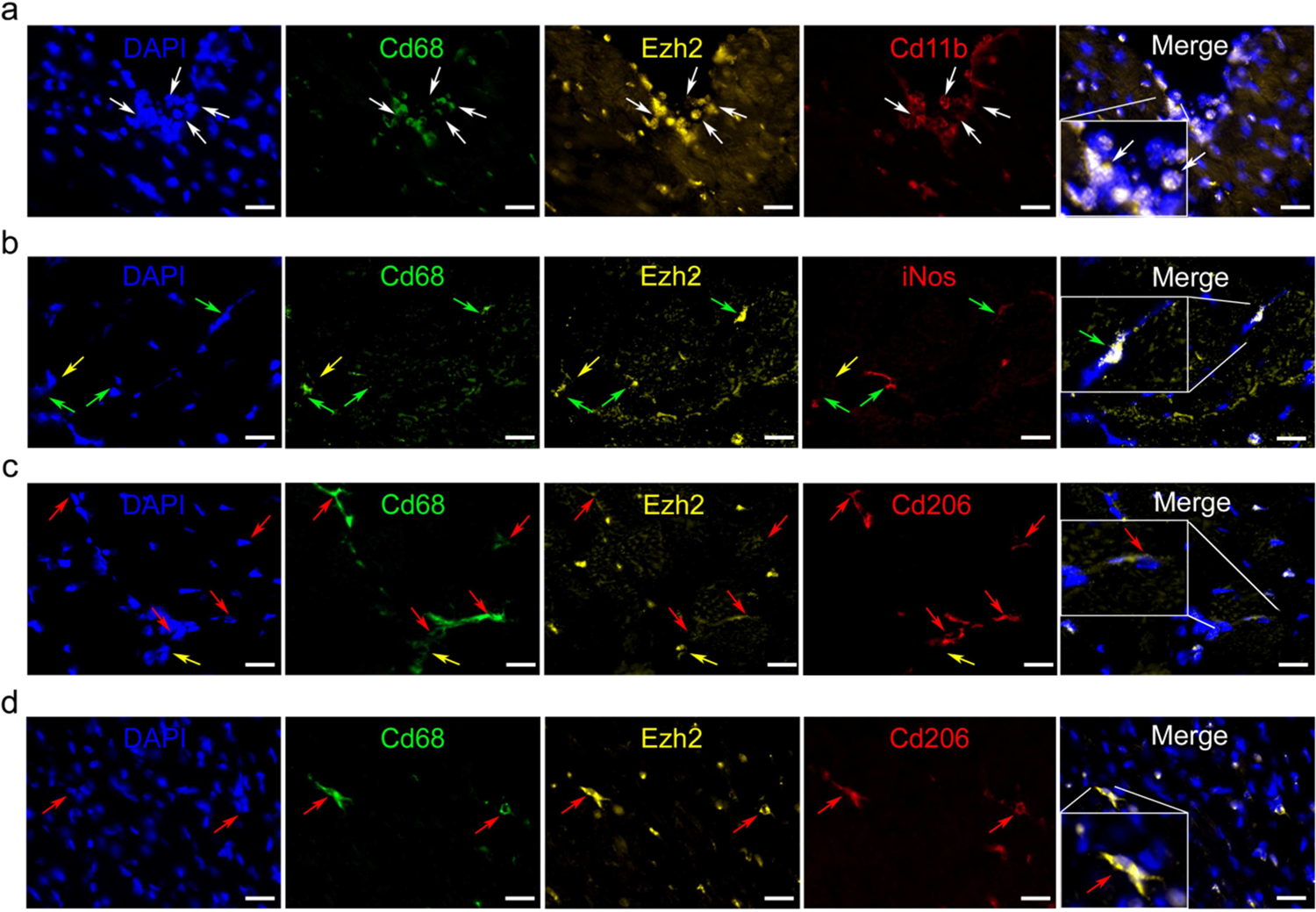
Ezh2 is translocated to the cytoplasm in cardiac M2 macrophages *in vivo* Representative pictures of cardiac immunostaining for Ezh2 in (a) Cd11b^+^/Cd68^-^ monocytes, (b) Cd68^+^/iNos^+^ M1 differentiated macrophages, (c) Cd68^+^/Cd206^+^ M2-like differentiated macrophages at 24h after coronary ligation in mice to induce MI or (d) cardiac-resident M2 macrophages in healthy sham mice. Nuclei were stained with DAPI (*blue*), myeloid markers were Cd11b (*red, panel a*), Cd68 (*green,* panels a-d). iNos (*red,* panel b), and Cd206 (*red*, panels c and d). Ezh2 (*yellow*) cellular localization was observed in each cell type but only appeared in M2 macrophages cytoplasm. Arrows indicate monocytes (white), non-determined macrophages (yellow), M1 (green) and M2 macrophages (red), scale bars represent 25 µm.

### Ezh2 cytoplasmic translocation promotes M2 macrophage polarization in vitro

To better understand the role of Ezh2 cytoplasmic translocation in M2-like cardiac macrophages post-MI, we reproduced myeloid cell differentiation *in vitro* starting from peripheral blood circulating monocytes to differentiated and mature polarized M1 or M2 macrophages. Peripheral blood circulating monocytes were subjected to negative magnetic selection before M-CSF-induced *in vitro* differentiation into non-polarized M0 macrophages followed by either LPS-induced M1 macrophage polarization or Il4 and Il10-induced M2 macrophage polarization. We confirmed our previous *in vivo* observations of a nuclear localization of Ezh2 in non-adherent (Fig. 2a) and adherent (Fig. 2b) monocytes, non-polarized M0 (Fig. 2c) and M1 (Fig. 2d) macrophages, while Ezh2 translocated to the cytoplasm in M2 macrophages (Fig. 2e). Interestingly, Ezh2 sub-cellular localization was not affected by classical or non-classical monocyte sub-population phenotype, as assessed by Ly6c expression (Fig. 2a), nor by selective M2a or M2c macrophage differentiation following single cytokine-induced polarization (Supplementary Fig. S1). Thus, this mechanism seems to be global rather than restricted a single monocyte or M2 macrophage sub-population phenotype. We hypothesized that Ezh2 exclusion from the nucleus may be critical to selectively allow M2 macrophage polarization and that it acts as an epigenetic checkpoint to regulate differentiation and polarization into immunoregulatory alternative M2 macrophages.

**Figure 2:**
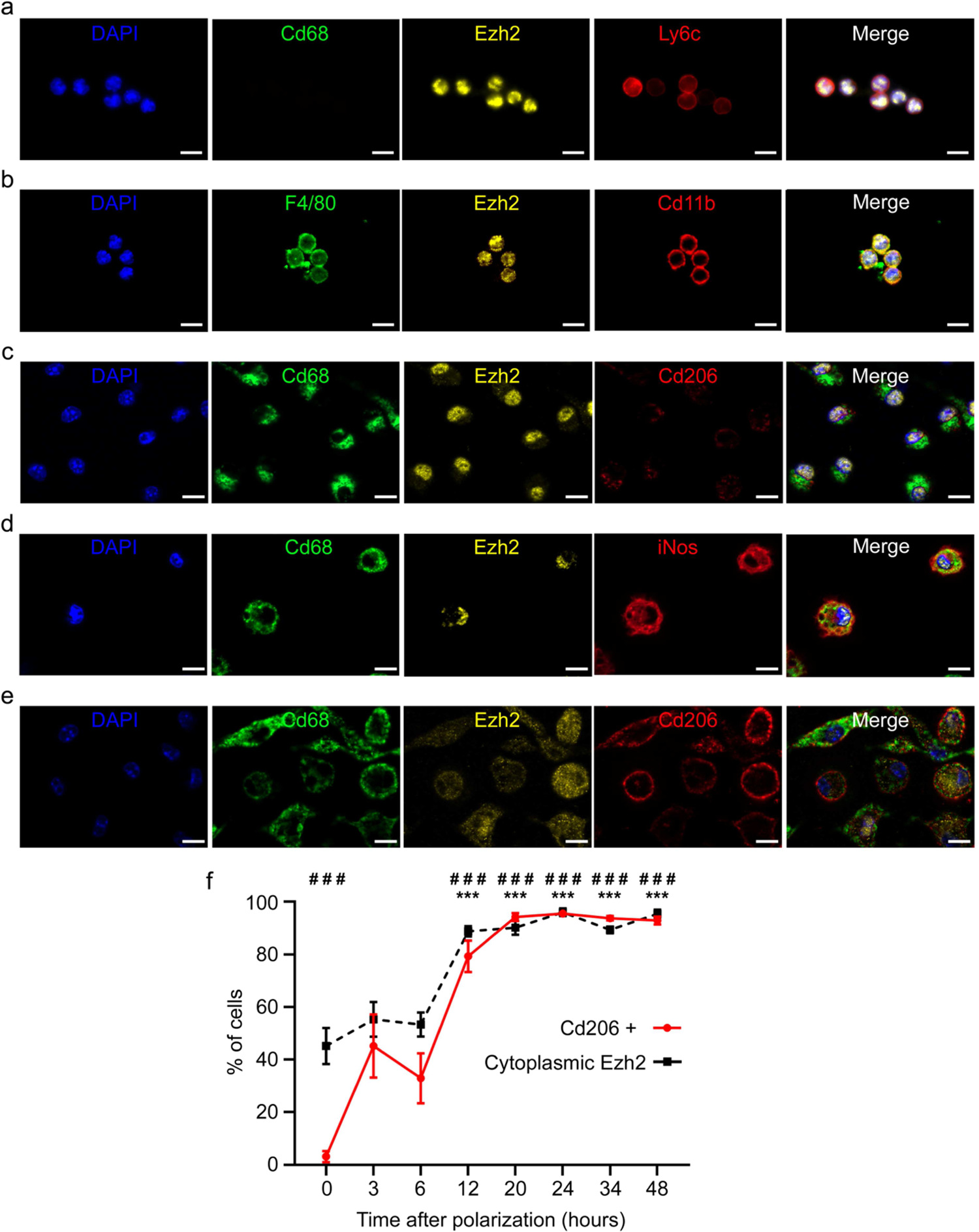
Ezh2 is translocated to the cytoplasm in myeloid cells during M2 polarization *in vitro* Representative pictures of immunostaining for Ezh2 in (a) non-adherent monocytes, (b) adherent monocytes, (c) non-polarized M0 macrophages, (d) M1 macrophages and (e) M2 macrophages, differentiated and polarized *in vitro*. Nuclei were stained with DAPI (blue), monocytes were identified based on Ly6c or Cd11b (*red*, panels a and b) expression, and macrophages based on Cd68 (*green, panels* a, and c-e) or F4/80 (*green*, panel b) expression. M1 or M2 macrophage phenotype was determined by expression of iNos (*red, panel d*) or Cd206 (*red,* panels c and e). Ezh2 (yellow) cellular localization was observed in each cell type. Scale bars represent 10 µm. Kinetics of Ezh2 subcellular localization and Cd206 expression (f) were assessed by immunostaining during M2 macrophage polarization. Data are represented as percentage of total cells mean values ± SEM of 3 independent experiments (n=3) each performed in duplicate for all time points. Asterisk (*) symbol indicates statistically significant difference between cells with either nuclear or cytoplasmic Ezh2 localization. Hashtag (#) depicts significant difference between Cd206 negative and positive cells. ### and ***p <0.001; Two-way ANOVA with Sidak’s multiple comparisons test.

To test this hypothesis, we induced macrophage differentiation *in vitro* as previously described, and examined Ezh2 cytoplasmic translocation as well as Cd206 expression kinetics. We observed an early Ezh2 nuclear to cytoplasmic translocation occurring within 3 hours after initiation of M2 macrophage polarization (Fig. 2f) prior to Cd206^-^ to Cd206^+^ switch, which appeared more than 6 hours after induction of polarization. We therefore conclude that Ezh2 might repress the M2 macrophage transcriptional programming during myeloid cell differentiation, which is relieved by nuclear export of Ezh2.

### M2 macrophage differentiation is epigenetically repressed by Ezh2 control of bivalent gene expression

Ezh2 acts as a transcriptional repressor of bivalent genes, characterized by the concomitant presence of both repressive Ezh2-dependent H3K27me3 and activating H3K4me3 epigenetic marks at their promoter regions^21^, ^22^. We decided to identify which bivalent genes may be possible direct targets of Ezh2 in macrophages. First, we exploited previously published ChIP-sequencing (ChIP-seq) data from CD14+ sorted human monocytes to establish a list of potential EZH2-regulated bivalent genes in human monocytes^23^, ^24^. Then, these putative EZH-2 regulated bivalent genes have been validated in CD14^+^sorted human monocytes by assessing the presence of H3K27me3 repressive mark found at the promoter of inactive genes, such as *PAX7* (Fig. 3a top panel), or H3K4me3 active mark at the promoter of active genes, such as *ELP3* (Fig. 3a second panel). Next, we established the full list of bivalent genes in CD14+ sorted human monocytes (supplementary table 1), which included *DLL1*, *VEGFA* and *IRF4* (Fig. 3a). Gene ontology (GO) analysis indicated that our list of bivalent genes in CD14+ sorted human monocytes are enriched in categories related to cardiac remodeling (e.g., cardiac right ventricle morphogenesis, heart development, ventricular septum morphogenesis), vascular remodeling (e.g., blood vessel development, angiogenesis), but also cell migration, wound healing, and cell fate commitment (Fig. 3a). This suggests that derepression, through EZH2 inactivation, of bivalent genes could promote cardiac, vascular and lymphatic repair functions in monocytes after MI. To investigate, we further examined the bivalent status of selected genes from our ChIP-qPCR analysis in our model of circulating monocytes isolated from human peripheral blood. As expected, we observed a significant enrichment of the H3K27me3 mark in the promoter region of *DLL1*, *VEGFA* and *IRF4* bivalent genes as well as in the promoter of *PAX7* inactive gene (Fig. 3c). Conversely, we found a significant enrichment of the activating H3K4me3 mark in the promoter of *DLL1*, *VEGFA* and *IRF4* bivalent genes as well as in the promoter of *ELP3* active gene. This data corroborates the bivalent status of *DLL1*, *VEGFA* and *IRF4* in human monocytes.

**Figure 3:**
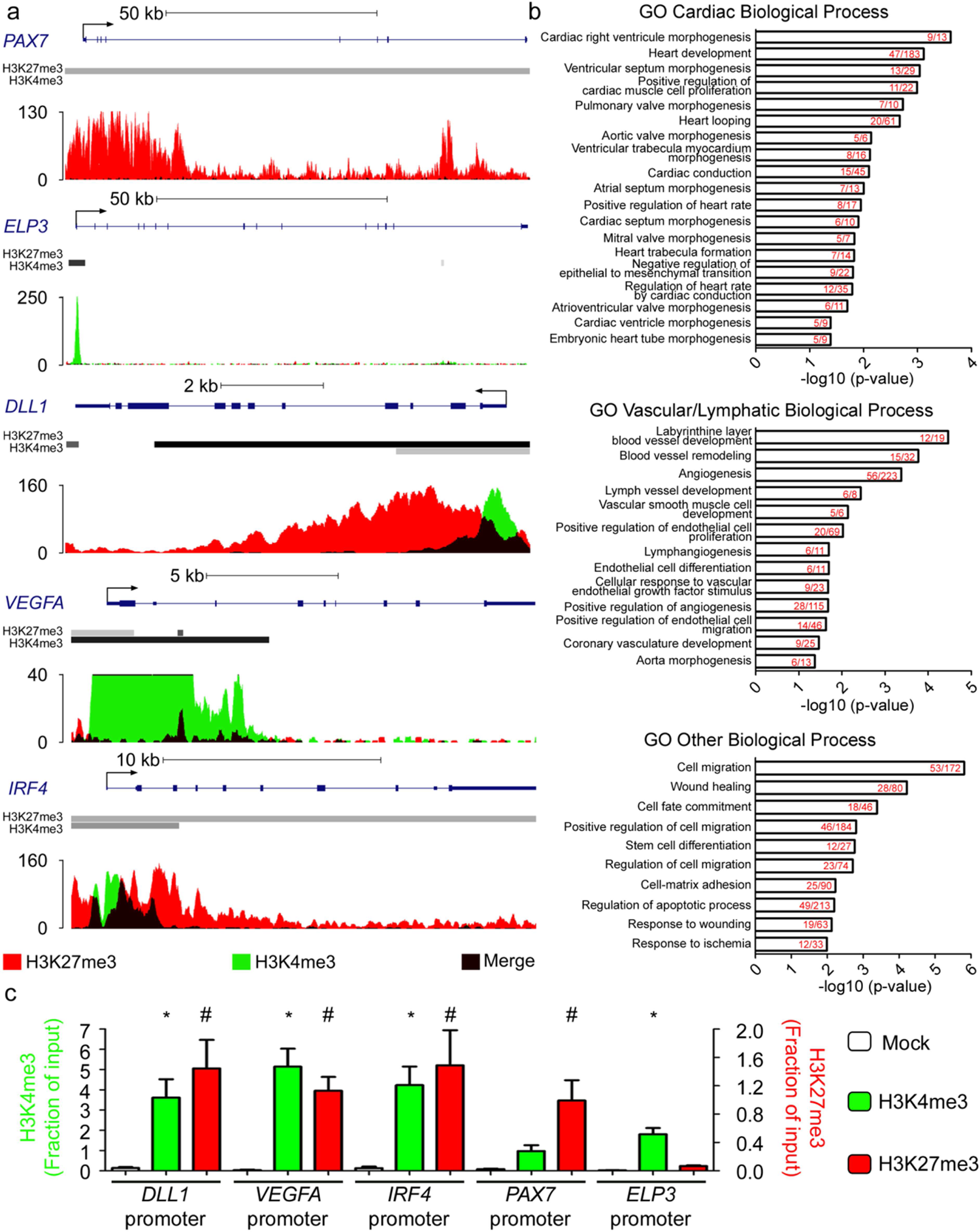
Ezh2-regulated bivalent genes in monocytes are implicated in cardiovascular repair processes Representative tracks and peak calling for promotors of inactive (*PAX7*), active (*ELP3*) and bivalent (*DLL1*, *VEGFA*, *IRF4*) genes (a) generated by ChIP-Seq analysis of human CD14+ monocytes. Representative Gene Ontology (GO) Biological Process categories significantly enriched for bivalent genes in human CD14+ monocytes. Number of genes revealed by the analysis within the number of overall genes in each GO biological process category indicated in red (b). Quantification of H3K27me3 inactive (red) versus H3K4me3 active (green) histone marks in the promoter regions of three bivalent and two non-bivalent genes, as assessed by ChIP-qPCR (c). Data represent mean fractions of input ± SEM of three independent experiments, corresponding to three different human donors, performed in duplicate (n=3). Asterisk (*) symbol indicates H3K4me3 statistically significant enrichment compared to the inactive *PAX7* promoter (Kruskal-Wallis test). Hashtag (#) depicts H3K27me3 significant enrichment compared to the active *ELP3*: *p <0.05 and # p <0.05.

### Pharmacological inhibition of EZH2 enhances bivalent gene expression to influence monocyte function in vitro

To determine the functional role of EZH2 on bivalent gene expression in monocytes, we used epigenetic pharmacological inhibition of EZH2 methyltransferase activity based on GSK-343 treatment. Once optimal inhibitory conditions (Supplementary Fig. S2A) and absence of nucleo-cytoplasmic translocation (Supplementary Fig. S2B) with GSK-343 were determined; we investigated by RNA-sequencing (RNA-seq) the transcriptomic impact of GSK-343 in isolated human monocytes. This experiment, performed using independent pools of selected peripheral blood monocytes from three distinct non-coronary patients, identified 85 statistically up-regulated genes and 48 statistically down-regulated genes as compared to vehicle control (Fig. 4a). GO analysis indicated that the 48 down-regulated genes are enriched in categories related to collagen binding, heart development, interleukin-1 binding, wound healing, and negative regulation of transcription (Supplementary Fig. S3A and supplementary table 2). In contrast, up-regulated genes are enriched in categories related to chemotaxis, response to hypoxia, angiogenesis and inflammatory response (Supplementary Fig. S3B, and supplementary table 3). The full lists of down- and up-regulated genes and GO categories are available respectively in tables 2 and 3.

**Figure 4:**
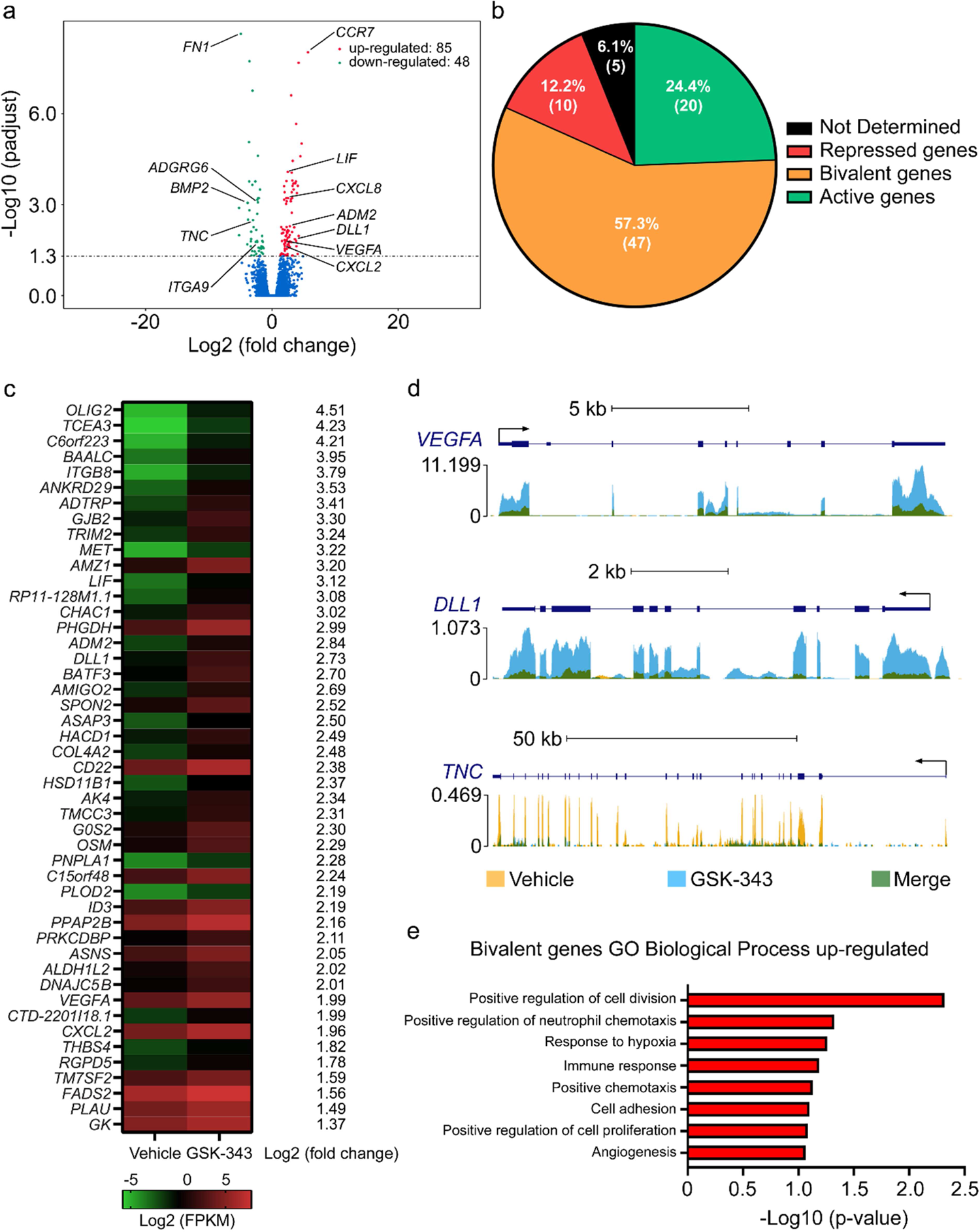
Ezh2 pharmacological inhibition with GSK-343 increases bivalent gene expression and enhances human monocyte homing and angiogenic functions *in vitro* Global changes in gene expression upon GSK-343 treatment versus vehicle treatment analyzed by mRNA-seq (a) in cultured human monocytes. Volcano plot shows upregulated (red) and downregulated (green) genes (combined data from three different donors, n = 3). Classification of GSK-343-induced genes according to promoter status (b). Heatmaps of all significantly up-regulated bivalent genes identified by RNA-seq analysis in human monocytes after GSK-343 treatment (c). Data obtained from three independent donors (n = 3) are expressed in log2 (FPKM). Representative tracks of changes, induced by GSK-343 treatment, in the epigenetic landscape in the promotor region of select up- and down-regulated genes in human monocytes (d). Representative Gene Ontology (GO) Biological Process categories significantly enriched for GSK-343-induced bivalent genes in human monocytes (e).

To get more insight into the molecular mechanisms and identify direct target genes of EZH2 modulation by GSK-343, we compared the list of bivalent genes identified in CD14+ human monocytes, established by ChIP-seq analysis, and the list of GSK-343-induced up-regulated genes, identified by RNA-Seq, in circulating human monocytes. We found that the majority of genes up-regulated following GSK-343 treatment were bivalent genes (47 out of 85 altered genes; 57.3 %). In contrast, the proportion of either active (20 out of 85 genes; 24.4 %) or repressed (10 out of 85 genes; 12.2 %) genes was minor, with some genes (5 out of 85 genes; 6.1 %) for which epigenetic status remains uncharacterized (Fig. 4b). Interestingly, within the full list of EZH2 direct target genes thus established (Fig. 4c), we found *VEGFA* (> 5-fold up-regulated in monocytes following GSK-343 treatment), a well-known master gene activator of angiogenesis and vasculogenesis and *DLL1* (> 6-fold up-regulated), a member of NOTCH signaling pathway regulating vascular morphogenesis and remodeling (Fig. 4d). Among the 48 down-regulated genes, we noted *TNC* (approximately 8-fold reduced as compared to vehicle treated monocytes), involved in cardiac fibrosis and acceleration of adverse ventricular remodeling after MI (Fig. 4d). Altogether our RNA-seq data identified a pool of 47 EZH2 bivalent gene targets (Fig. 4b and c) implicated in chemotaxis (*VEGFA*, *MET, THBS4* and *CXCL2*), response to hypoxia (*PLOD2*, *VEGFA* and *PLAU*), immune response (*LIF*, *CD22, OSM* and *CXCL2*) and angiogenesis (*VEGFA, ADM2, DLL1,* and *COL4A2*) (Fig. 4e) suggesting that EZH2 inhibition with GSK-343 may promote monocyte cardiac repair function.

### EZH2 pharmacological inhibition specifically resolves H3K27me3 level at bivalent gene promoters to activate transcription

To verify that the upregulated expression of bivalent genes in response to GSK-343 treatment was directly due to alleviation of EZH2 gene repression we next investigated the level of the repressive H3K27me3 chromatin mark in the promotor regions of selected target genes. Indeed, we found, using ChIP-qPCR analysis, that *in vitro* treatment of human monocytes for 72h with GSK-343 decreased H3K27me3 levels at *DLL1*, *VEGFA* and *IRF4* gene promoters (Fig. 5a). Interestingly, H3k27me3 levels were not modified at inactive gene promoters, such as *PAX7,* nor in constitutive active gene promoters, such as *ELP3* (Fig. 5a). As expected, the levels of the H3K4me3 active mark were not modified upon GSK-343 treatment in the promotor regions of either bivalent genes (e.g. *DLL1*, *VEGFA* and *IRF4)* or active genes (*ELP3*). This epigenetic activating mark was also not detectable at inactive gene promoters (*PAX7*), in line with suppressed transcription activity in monocytes (Fig. 5b). Taken together, these results demonstrate the specificity of pharmacological inhibition of EZH2 as an approach to selectively remove epigenetic gene repressive marks present on bivalent gene promoters, without affecting their activating epigenetic marks.

**Figure 5:**
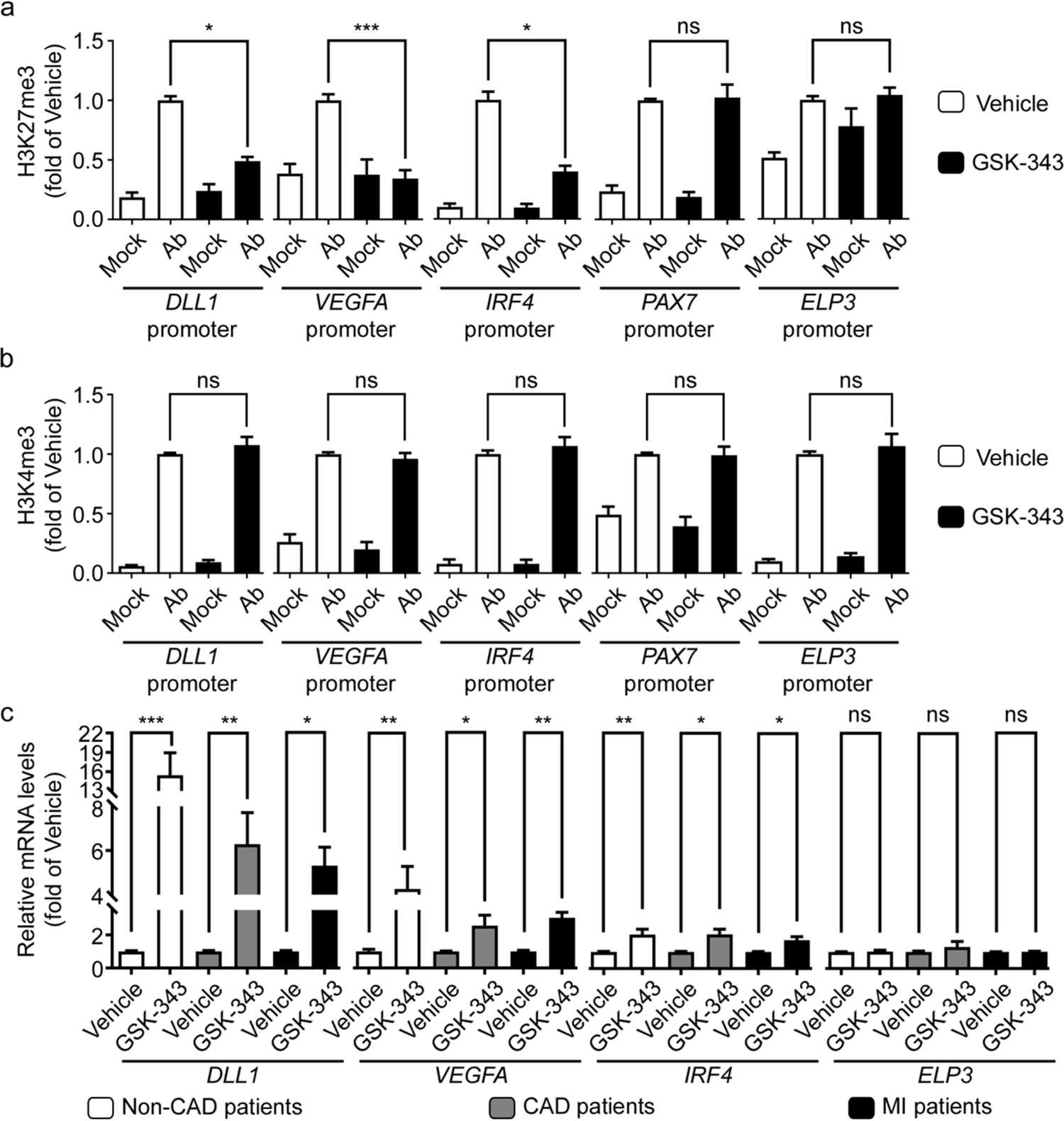
Pharmacologic inhibition of EZH2 increases specifically bivalent gene expression through H3K27me3 demethylation in monocytes *in vitro* Enrichment of selected bivalent (*DLL1, VEGFA, IRF4*), inactive (*PAX7*) and active (ELP3) gene promoters in H3K27me3 (a) and H3K4me3 (b) as assessed by ChIP-qPCR. Ab indicates the enrichment obtain with either H3K27me3 (a) or H3K4me3 (b) targeting antibodies, while Mock represents the data obtain after immunoprecipitation with control IgG. Data are represented as mean fractions of input normalized to vehicle-treated monocytes ± SEM of three independent experiments corresponding to three different donors performed in duplicate (n=3). The p values are depicted as asterisks in the figures as follows: ***p <0.001; **p <0.01; *p <0.05; ns: non-significant enrichment compared to vehicle-treated monocytes. Transcript levels of indicated genes were measured by RT-qPCR following treatment with vehicle or GSK-343 of selected monocytes from non-coronary patients, CAD or AMI patients (c). RT-qPCR values are expressed as mean percentages of vehicle-treated monocytes ± SEM with B2M serving as internal control of four independent experiments corresponding to four different donors per group performed in duplicate (n=4). ***p <0.001; **p <0.01; *p <0.05; ns: non-significant.

To gain further insight into the potential therapeutic applications of EZH2 inhibition in human monocytes after MI, we isolated circulating monocytes from patients admitted to the department of cardiology of Rouen university hospital. Patients were divided into three groups corresponding to 1) patients without coronary disease; 2) patients diagnosed with stable coronary artery disease (CAD); and 3) patients admitted after acute MI (AMI). Patient characteristics are detailed in supplementary table 4. In line with previous findings, we observed a slight but non-significant increase in circulating monocytes levels after MI, as compared to non-coronary patients (Supplementary Fig. S4). Monocytes collected from patients of the three different groups were then treated with GSK-343, as described above, followed by RT-qPCR analysis of gene expression of selected bivalent (*DLL1*, *VEGFA*, *IRF4*) or active (*ELP3*) genes to assess GSK-343 treatment effects. Interestingly, we observed a significant up-regulation of all selected bivalent genes in all three patient groups, while the expression of constitutive active genes remained unaltered (Fig. 5c). These results demonstrate that GSK-343-mediated EZH2 inhibition is efficient to activate bivalent gene expression in human monocytes irrespective of the underlying disease conditions. Other genes modified expressions identified by RNA-seq were confirmed by RT-qPCR including *ADM2*, *BMP2*, *CXCL2*, *FN1* and *TNC* (Supplementary Fig. S5). Moreover, a targeted transcriptomic array of a murine monocyte cell line, focused on assessment of genes selectively expressed in different myeloid cell lineages, revealed that treatment with GSK-343 induces an expression shift similar as observed in M2 macrophage profile (Supplementary Fig. S6A). These results of GSK-343 were confirmed in both mouse (Supplementary Fig. S6B and C) and human (Supplementary Fig. S6D and E) freshly-isolated monocytes as compared with *in vitro* differentiated M0, M1 or M2 macrophage subtypes. Altogether our *in vitro* data show that EZH2 acts as an epigenetic check point inhibiting, at the chromatin level, bivalent gene expression to prevent monocyte/macrophage differentiation into an M2-like immunomodulatory cell type. Conversely, treatment of monocytes with a pharmacologic inhibitor of EZH2 activity favored suppression of the H3K27me3 repressive epigenetic mark at the promoter of bivalent genes, thus enhancing their transcription to initiate a cell differentiation program and/or a cell fate commitment into M2-like immunoregulatory macrophages. These observations argue for a potential use of GSK-343 as a novel epigenetic treatment to accelerate inflammatory resolution after MI to limit cardiac dysfunction and heart failure development.

### GSK-343 accelerates inflammation resolution and prevents subsequent cardiac dysfunction development in mouse model of MI

Next, we evaluated the effects of GSK-343 treatment on cardiac inflammation and cardiac function in a mouse model of MI induced by permanent left coronary ligation. As GSK-343 is a poorly soluble epigenetic drug, we used cyclodextrin-based Captisol^®^, as previously described for daily intraperitoneal injections^25^. Because our *in vitro* data indicated that EZH2 inhibition promotes immunomodulatory functions in monocytes, we first examined circulating and cardiac immune cell levels post-MI using flow cytometry. In line with previous reports, circulating as well as cardiac classical pro-inflammatory monocyte populations were found to be increased at 3 days after MI. Interestingly, GSK-343 treatment significantly reduced classical blood monocyte (Ly6C^hi^CXCR3^hi^)/inflammatory (M1)-like levels while increasing cardiac levels of the same population in both infarct scar and border zone at 3 days post-MI (Fig. 6a, left and middle panels). This indicates that Ezh2 inhibition may accelerate cardiac inflammatory kinetics after MI. In agreement, we found that at 8 days after MI the non-classical immunomodulatory monocyte (Ly6C^lo^CXCR3^hi^)/M2-like population was significantly increased in both infarct and Border Zone (BZ) areas in GSK-343-treated as compared to vehicle-treated mice (Fig. 6a, right panel). Notably, the levels of these alternative monocytes in GSK-343-treated mice were similar as in sham controls, arguing for accelerated resolution of inflammation with GSK-343. Similarly, cardiac macrophage levels tended to be more rapidly increased in GSK-343-treated mice as compared to vehicle controls (Supplementary table 5). To further characterize the cardiac inflammatory response, we assessed cardiac inflammatory cytokine expression by RT-qPCR. In support of faster resolution of inflammation upon GSK-343 treatment, we observed significantly reduced expression of pro-inflammatory cytokines *Il1b* and *Il6,* 8 days after MI in hearts, as compared to vehicle-treated mice (Fig. 6b). For example, the MI-induced increase in cardiac *Ccl2* (a.k.a monocyte chemoattractant protein) expression, was significantly decreased in GSK343-treated mice, while Ccl21 was increased, suggesting reduced monocyte recruitment and/or increased immune cell efflux via lymphatics in GSK-343 treated mice (Fig. 6b). Interestingly, we did not observe any modification of this gene expression 3 days post-MI (supplementary table 6).

**Figure 6:**
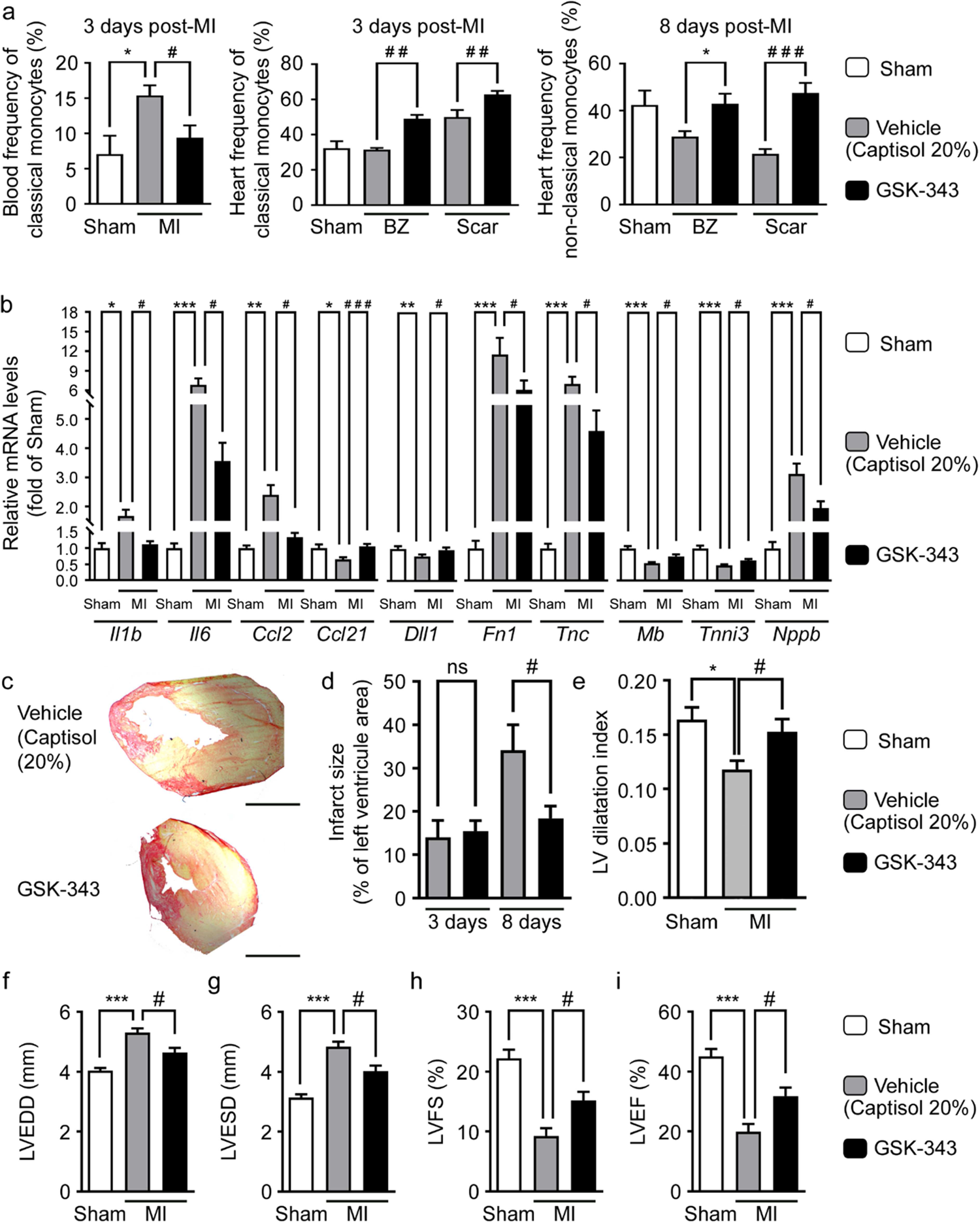
Pharmacological Ezh2 inhibition with GSK-343 accelerates cardiac inflammatory resolution and prevents infarct expansion and subsequent cardiac dysfunction after MI Circulating (3 days after MI) and cardiac (3 days and 8 days after MI respectively for center and right panels) classical (Cd11b^hi^Ly6c^hi^) and non-classical (Cd11b^hi^Ly6c^lo^) monocyte population frequencies were evaluated by flow cytometry. Samples were collected from sham (n=4) and MI mice treated daily either with vehicle (captisol 20%, n=13 blood samples, n=4 cardiac samples) or GSK-343 either from the infarcted scar area or from the healthy border zone (BZ). (a). Data are expressed as mean frequency ± SEM of live CD45^+^/Cd19^neg^ cells. Cardiac transcript levels of indicated genes measured by RT-qPCR from sham (n=8), or vehicle (n=16) or GSK-343-treated (n=18) mice at 8 days after MI (b). Values expressed as mean percentages of sham ± SEM with B2m serving as internal control. Representative pictures of infarct scar and M-mode echocardiography in sham, vehicle and GSK-343 treated mice at 8 days post-MI (c). Scale bar represents 1 mm. Infarct size assessed in vehicle (n=5-7) or GSK-343 treated (n=6-7) MI-mice (d). Data represent mean percentage of left ventricle area ± SEM with # p <0.05; ns: non-significant. Quantitative analysis of LV dilatation index (e) and echo parameters for wall thickness (e), LVEDD (f), LVESD (g), LVFS (h) and LVEF (i) in sham (n=8), vehicle (n=14) and GSK-343 (n=18) treated mice at 8 days post-MI. Data are presented as means ± SEM. Asterisk (*) and hashtag (#) symbols indicates statistically significant difference compared to sham and vehicle condition respectively after Kruskal-Wallis test. # p <0.05, *** p <0.001.

Altogether, our data argue for accelerated resolution of the inflammatory phase post-MI with GSK-343 treatment. Further, as expected from RNA-seq data in monocytes, cardiac *Dll1* bivalent gene expression was restored with epigenetic drug treatment while the expression of both *Fn1* and *Tnc* pro-fibrotic genes was significantly reduced as compared to vehicle treated animals (Fig. 6b). However, different from *in vitro* data, we did not observe a significant increase in cardiac *Vegfa* expression upon GSK-343 treatment (supplementary table 6), although we identified it as a bivalent gene. This suggests that recruited myeloïd cells might not be the principal source of *Vegfa* during early steps of cardiac repair after MI. In accord with absence of major angiogenic or lymphangiogenic effects, both cardiac vessel density and lymphatic densities were not significantly increased after 8 days in GSK-343-treated groups as compared to vehicle controls (Supplementary table 5).

By accelerating the resolution of the inflammatory phase, GSK-343 treatment may influence cardiac scar formation strength or kinetics. To investigate this, we evaluated cardiac function and morphology after MI. Although infarct size was similar 3 days after MI, we noticed a stabilization of infarct scar size, with less expansion by 8 days after MI in GSK-343 as compared to vehicle treated groups (Fig. 6d and e). In agreement with less cardiac stretching due to infarct expansion, GSK-343-treated mice displayed reduced cardiomyocyte hypertrophy at 8 days post-MI (Supplementary table 5). Consistent with these findings we observed higher expression 8 days after MI of both *Mb* and *Tnni3*, encoding respectively myoglobin and troponin I cardiomyocyte-specific genes in GSK-343-treated mice suggesting limitation of cardiomyocyte cell death or dysfunction in the border zone (Fig. 6b). We conclude that Ezh2 inhibition with GSK-343 prevents infarct scar expansion through expedited inflammatory phase resolution, leading to increased scar maturation and potentially reduced cardiomyocyte remodeling or dysfunction. Indeed, we observed partial prevention of cardiac dysfunction 8 days post-MI, with reduced left ventricular (LV) dilation, including improvement of both diastolic (Fig. 6f) and systolic (Fig. 6g) diameters, leading to rescue of LV fractional shortening (Fig. 6h) as well as ejection fraction (Fig. 6i) after GSK-343 daily treatment. This is further confirmed by our observation of significantly reduced expression of *Nppb*, a well-known biomarker of deleterious cardiac remodeling and heart failure at 8 days post-MI. However, none of these parameters were improved by GSK-343 at 3 days post-MI (supplementary table 7). We conclude that GSK-343 treatment accelerated inflammatory resolution, leading to improved infarct scar maturation, cardiomyocyte protection and subsequent reduction of infarct expansion, resulting in reduced cardiac dysfunction after MI.

## Discussion

We report, to the best of our knowledge, for the first time a mechanism involving cytoplasmic translocation of EZH2 as a cellular epigenetic switch to facilitate M2 macrophage polarization linked to derepressed expression of bivalent genes. This finding of cytoplasmic translocation of EZH2, while unreported in macrophages, has previously been described in T cells during actin polymerization, where EZH2 was found to interact with Vav1 to regulate T-cell-receptor-mediated signalling^26^.

Moreover, during megakaryopoiesis, Notch1 was described to induce EZH2 cytoplasmic translocation where it interacts with LIM domain kinase-1 (LIMK1), resulting in reduced cofilin phosphorylation. This increases cofilin activity and subsequently decreases filamentous actin content, preparing the cells for cell shape and size modifications required for megakaryocyte formation^27^. Finally, a recent report has indicated another cytoplasmic role for EZH2 linked to breast cancer metastasis. It was demonstrated that p38-induced phosphorylation of EZH2 at Tyr_367_ promoted its cytoplasmic retention in breast cancer cells. This in turn enhanced EZH2 binding to vinculin and other cytoskeletal regulators, leading to promotion of cell adhesion, migration, invasion and subsequent development of breast cancer metastasis^28^. These data depict novel cytoplasmic roles of EZH2, which may also relate to our transcriptomic findings of improved homing and chemotaxis/ migration of M2 macrophages and GSK-343 treated monocytes. Interestingly, besides LIMK1 and Vav1, additional cellular partners of EZH2 have been identified and may be involved in cytoplasmic translocation of EZH2 during M2 macrophage polarization. Among these putative partners, long non-coding RNA (lnc RNA) are interesting targets.

Indeed, EZH2 RNA Immune-Precipitation-sequencing (RIP-seq) experiments to identify tissue-specific lnc RNA partners of EZH2 confirmed interaction with both Cardiac Hypertrophy Associated Epigenetics Regulator (CHAER)^29^, ^30^ and HOx Transcript Antisense RNA (HOTAIR) lnc RNA in both heart and blood^29^. Whereas Chaer was found to directly interact with PRC2, leading to inhibition of Ezh2-mediated repression of hypertrophic genes in the heart^30^, Hotair interacted with PRC2 and enhanced EZH2 activity, leading to increased myofibroblast differentiation by promoting collagen and α-SMA expression. It remains to be determined whether these partners could be involved in regulation of Ezh2 cytoplasmic translocation during M2 macrophage differentiation/polarization.

Although the data available on the role of EZH2 in the innate immune response remains scarce, a recent report examined the effects of Ezh2 deletion in macrophages and microglia. The authors reported that Ezh2 is required to repress Socs3 expression, necessary to promote pro-inflammatory gene expression, in response to Toll-Like Receptor (TLR)-mediated signaling during macrophage and microglia activation in autoimmune inflammation^20^. This is in agreement with our data staring EZH2 as a crucial epigenetic check point regulator that prevents M2 differentiation/polarization thus extending the duration and/or severity of cardiac inflammation after MI. In this study, we show for the first time that pharmacological inhibition of EZH2 with GSK-343 promotes the capacity of human monocytes to accelerate cardiac inflammatory resolution through derepression of bivalent genes Further, we propose that this epigenetic control of bivalent gene expression mediated by H3K27me3 demethylation by epigenetic enzymes such as JMJD3 and UTX. Indeed, previous studies have shown that IL4 up-regulates JMJD3 expression and activity in human monocytes. This in turn, decreases H3K27me3 level at *IRF4* promoter, increases RNA polymerase II recruitment and promotes subsequent *IRF4* mRNA expression. The implication of JMJD3 in bivalent gene reexpression was further confirmed by the inhibition of *IRF4* expression after treatment with a pharmacologic inhibitor of JMJD3, GSK-J4^12^. In our study, we observed a similar phenomenon (decrease of H3K27me3 level at *IRF4* promoter enhancing its mRNA expression) after treatment of human monocytes with the EZH2inhibitor GSK-343. This suggest that the balance between EZH2 and JMJD3 might be responsible for determining bivalent gene (*IRF4*, *VEGFA* and *DLL1*) promoter epigenetic mark H3K27me3 levels, which directly determines gene expression activity in monocytes during macrophage differentiation. Among these bivalent genes, *DLL1* expression by endothelial cells has been shown to promote conversion of Ly6C^hi^ classical monocytes into Ly6C^lo^ alternative monocytes *in vivo* and *in vitro*^31^. We propose that *DLL1* up-regulation in monocytes could participate in cell-cell communication through NOTCH2 activation to stimulate inflammatory-to-immunoregulatory monocyte switch. We postulate that EZH2 inhibition of M2 macrophage polarization is a global phenomenon, as we reproduced similar data from peripheral blood monocytes isolated from non-coronary as well as coronary and AMI patients. We also provide, for the first time, *in vivo* proof of the protective effects induced by pharmacological EZH2 inhibition in a mouse MI model with permanent coronary artery ligation. Our data are in agreement with a recent study highlighting EZH2 overexpression and cardiac enrichment of known gene targets of EZH2, such as *KLF15*, in ischemic cardiomyopathy patients. Consequently, EZH2 was proposed as a likely transcriptional regulator of gene expression in ischemic cardiomyopathy^32^. In our mouse MI model, we observed accelerated inflammatory resolution upon EZH2 inhibition, in agreement with our *in vitro* data on altered gene expression and polarization of monocytes. Of note, we observed decreased cardiac expression upon GSK-343 treatment of both *Il1b* and *Il6* pro-inflammatory cytokines and *Ccl2,* implicated in cardiac monocyte recruitment and early immune response, as well as restored expression of *Ccl21,* involved in late immune response and cardiac repair^33^. We speculate that restored *Ccl21* expression may relate to protection of cardiac lymphatics, as they constitute the main cardiac source of *Ccl21.* Moreover, our RNA-Seq data in human monocytes revealed that the CCL21 receptor, *CCR7*, is the highest up-regulated gene after GSK-343 treatment. This indirectly suggests that GSK-343 may have enhanced both cardiac and lymphatic homing of myeloid cells, *via* promotion of Ccr7/Ccl21 signaling, resulting in accelerated clearance of cellular debris and inflammation in the heart^34^ leading to prevention of infarct scar expansion. Further, GSK-343 treatment may also have directly altered fibrosis and scar maturation, as suggested by the reduction in *Fn1* and *TnC* expression in the heart.

However, in addition to modulating collagen production in part by enhancing TGFβ signaling, TnC has also been shown to act as a trigger for monocyte/macrophage recruitment^35^ leading to accelerated adverse ventricular remodeling after MI^36^ and heart failure development. The immunomodulatory effects of TnC seem to involve suppression of *Irf4* expression, necessary for M2 macrophage polarization. Taken together, our findings suggest that GSK-343-induced *Irf4* up-regulation in monocytes together with cardiac *TnC* down-regulation could potentiate M2 macrophage polarization to accelerate cardiac repair and limit inflammation. In conclusion, our study brings, for the first time, evidence of the key role of EZH2 as an epigenetic check-point regulator that prevents M2 macrophage polarization.

Promisingly, pharmacological EZH2 inhibition derepresses specific bivalent genes, resulting in increased expression of genes favoring M2 polarization. In the setting of MI, the enhanced cardiac recruitment and activity of non-classical monocytes induced by EZH2 inhibitions resulted in accelerated inflammatory resolution and decreased infarct scar expansion, leading to a reduction of cardiac remodeling and dysfunction after MI. In conclusion, our data suggest EZH2 as an attractive novel therapeutic target to reduce cardiac inflammation and limit heart failure development following MI.

## Online methods

### Human study design and approvals

Between Nov 2018 and June 2020, the EPICAM prospective study enrolled 48 patients at Rouen University Hospital. The CPP Ile de France V has approved the study august 10th 2018 (RCB: 2018-A02108-47). All patients read information sheet and accepted to participate to the study before blood collection. Inclusion criteria were: 1) age greater than or equal to 18 years and 2) admission to hospital for a coronary angiography. The exclusion criteria were 1) Infectious diseases, 2) Current pregnancy or breastfeeding, 3) Obesity (BMI > 30 kg/m²), 4) Haematological pathologies, 5) Anaemia and 6) Inflammatory and autoimmune pathologies. During the coronarography procedure, peripheral blood was collected into 4 BD Vacutainer® EDTA K2 tubes (Becton Dickinson Cat#367862). In-hospital data were entered into a dedicated database.

### Animal study design and approvals

All animal experiments performed in this study were approved by the regional ethics review board in line with E.U and French legislation (01181.01 / APAFIS #8157-2016121311094625-v5 Normandy). We only used female mice for MI studies as they display lower mortality than male in the MI model, which help us reduce the numbers of animals included in our studies. In addition, females express both alleles of the H3K27me3 demethylase *Kdm6a* gene encoding the Utx protein. Mouse peripheral blood samples were collected in BD Vacutainer® EDTA K2 tubes (Becton Dickinson Cat#367862) from C57Bl/6J female mice aorta.

### Cell isolation and culture

Monocytes were directly or indirectly (after red blood cell lysis (eBioscience Cat#00-4333-57) isolated by negative selection using EasySep™ monocyte isolation kits for human (StemCell Technologies Cat#19669) and mouse (StemCell Technologies Cat#19861) cells from fresh peripheral blood samples according to the manufacturer’s instructions. TIB-204™ (WEHI-265.1) mouse monocyte cell line (ATCC Lot#4249478) was used in this study. All monocytes were cultured in Dulbecco’s modified Eagle’s medium (DMEM, Gibco, Cat# 41966-029) supplemented with 50 µM of 2-mercaptoethanol (Gibco Cat#21985-023), 10% Fetal Bovine Serum (FBS, Gibco Cat#10500-064) and Penicillin/Streptomycin (Sigma-Aldrich Cat#P4333) seeded at a cellular density > 150 000 ¢/cm² for primary monocytes and > 200 000 ¢/ml for TIB-204. M0 macrophages were differentiated from TIB-204 with 100 µM Phorbol 12-myristate 13-acetate (PMA, Sigma-Aldrich # P8139-1MG) and from selected primary monocytes using 50 ng/ml of either murine or human Macrophage-Colony Stimulating Factor (M-CSF) for 4 days. Further differentiation of M0 macrophages into polarized macrophages was obtained with 50 ng/ml LipoPolySaccharide (LPS, Sigma-Aldrich Cat#L6529) for M1 and with a combination of both IL4 and IL10 at a final concentration of 20 ng/ml respectively for M2 polarized macrophages for 2 days.

### Antibodies and reagents

For differentiation of primary monocytes into M0 macrophage *in vitro*, murine (Peprotech Cat#315-02) or human (StemCell Technologies #78057.1) M-CSF was used. The subsequent M0 into M2 macrophage polarization was obtained using a cocktail of murine Il4 (Peprotech Cat#214-14) and Il10 (Peprotech Cat#210-10) or human IL4 (StemCell Technologies Cat#78045.1) and IL10 (StemCell Technologies Cat#78024.1). M2 macrophage subpopulation phenotypes were obtained after incubation with 20 ng/ml of murine Il4 (Peprotech Cat#214-14), Il10 (Peprotech Cat#210-10) and Tgfb1 (R&D Sytems Cat#7666-MB-005) alone or in combination depending on the M2 macrophage subtype desired. The Ezh2 inhibitor GSK-343 (Sigma-Aldrich Cat#SML0766) was dissolved in dimethyl sulfoxide (DMSO, Sigma-Aldrich Cat#D1435) and used at a final concentration of 5 µM for *in vitro* experiments.

The following antibodies were used at the indicated dilutions: Immunohistochemistry: rat anti-CD11b (BD Pharmingen Cat#557395, 1:1500), rat anti-CD68 (eBiosicence Cat#14-0681, 1:800), goat anti-CD206 (Thermo Fisher Cat#PA5-46994, 1:1500), rabbit anti-EZH2 (D2C9, Cell Signaling Cat#5246, 1:1000), rat anti-iNOS (W16030C, Biolegend Cat#696802, 1:500), WGA-FITC (Interchim#FP-CE8070, 1:100), biotinylated rabbit anti-Lyve1 (eBioscience#13-0443, 1:400), biotinylated rat anti-CD31 (BD pharmingen#553371, 1:50); biotinylated rabbit anti-Cx3cr1 (BIOSS Cat#bs-1728R-Biotin, 1:100), and rat anti-Ly6c (W16030C, Abcam Cat#ab15627, 1:400).

Chromatin Immunoprecipitation: 2 µg of the following antibodies were used: rabbit anti-H3K4me3 (Millipore Cat#07-473), rabbit anti-H3K27me3 (Millipore Cat#07-449), or normal rabbit IgG (Millipore Cat#12-370). Flow cytometry (FACS): anti-mouse CD45 (30F11); anti-mouse CD11b (M1/70); anti-mouse Ly6C (HK1-4); anti-mouse CX3CR1 (SA011F11); anti-mouse CD86 (GL-1); anti-mouse CD3 (17A2); anti-mouse CD115 (AFS798); anti-mouse CD11c (N418); anti-mouse CD19 (6D5); anti-mouse CD206 (C068C2); anti-mouse IA-IEk (MHCII) (M5/114.15.2); anti-mouse F4-80(BM8) (All from Biolegend, used between 1/200 to 1/400), LIVE/DEAD Viability/Cytotoxicity (Invitrogen, 1/1000) Western blotting: anti-pan histone H3 (Millipore Cat#07-690, 1:100 000), anti-H3K27me3 (Millipore Cat#07-449, 1:2000).

### Mouse model of Myocardial infarction

Left ventricular (LV) MI was induced in 20-22g C57Bl/6 female mice by permanent ligation of the left descending coronary artery as previously described^37^. GSK-343 (20 mg/kg, Interchim Cat# XLR94D),was dissolved in cyclodextrin-based Captisol® as a solubilization vehicle at 20% final concentration, as advised by the manufacture, to perform daily intraperitoneal injections 6 hours after surgery and daily until 3 to 7 days post-MI. In controls vehicle solution (Captisol® 20%) was administered by intraperitoneal injection.

### Immunohistochemistry analysis

Frozen OCT-embedded mouse hearts were sectioned on a cryostat with a section size of 8 µm. Air dried sections were fixed in ice cold acetone (−20°C) for 10 min before a 1h permeabilization step in PBS containing 0.1% saponin; 1% BSA; 4% donkey serum. Incubation with desired primary antibody was performed 1h to overnight. Primary antibody signal was revealed using donkey anti-specie fluorescent antibody. Pictures were taken on Zeiss microscope imager Z.1 equipped with an apotome.

### Immunocytochemistry analysis

TIB-204 and primary mouse monocytes were stained in suspension while adherent macrophages were directly stained on 14 mm ø coverslips. Briefly, cells were fixed for 10 min in 1% or 4% paraformaldehyde respectively for suspension or adherent cells. Permeabilization was then performed using PBS containing 0.25% Triton X-100 or 0.1% saponin respectively for suspension or adherent cells. Cells were incubated overnight with desired primary antibodies, as described above, at indicated concentration after 30 min incubation in PBS containing 0.1% saponin; 1% BSA; 4% donkey serum.

### RNA extraction

Total RNA was extracted from at least 3×10^6^ human and 5×10^5^ murine monocytes treated or not with 5 µM GSK-343 for 72h using NucleoSpin RNA XS kit (Macherley-Nagel Cat# 740902) for *in vitro* studies or from 5 mg of cardiac tissue using RNeasy Mini Kit (Qiagen Cat#74104) including a genomic DNA digestion step with DNAse I (Qiagen Cat#79254).

### Gene expression profiling by RNA-Sequencing

For RNAseq, we used total RNA extracted from 3 independent non-coronary patients treated with 5 µM GSK-343 or vehicle for 72 h therefore representing three biological replicates. Total RNA was sent for library preparation, mRNA sequencing and bio-informatics analysis to Novogene Company (Novogene company Ltd, Cambridge, UK). 400 ng of total RNA with a concentration > 20 ng/ml and a RIN ratio > 8 were used for Poly-A mRNA-seq, sequencing length of 150 nt paired-end (PE150) with the Illumina NovaSeq 6000. Reads were aligned to the human genome build hg38 and the transcript assembly GRCh38 from Ensmbl using Hisat2 version 2.0.5^38^.

Transcript quantification was performed with featureCounts from the subread package^39^. Differential analysis, including normalization, was performed with DESeq2 bioconductor package version 1.22.1^40^ using R version 3.5.1. Genes significantly up- or down-regulated upon epigenetic combined drug treatment (adjusted p-value < 0.05) were submitted to DAVID Gene Ontology Analysis version 6.7^41^, ^42^. To generate heat maps, expression values in FPKM for genes within the selected GO Categories were retrieved. Heat maps were generated with Cluster version 3.0^43^ and JavaTree View version 1.1.6r4^44^.

RNA-Seq data are available into the public GEO database (GSE165543) with the following protected access for reviewers using the provided link and token: Link: https://www.ncbi.nlm.nih.gov/geo/query/acc.cgi?acc=GSE165543 Token: mvmvwsogdhszlez

### Reverse Transcription-quantitative Polymerase Chain Reaction (RT-qPCR)

RT-qPCR was performed with 0.4 to 1.5 µg total RNA isolated as previously described. Extracted RNA was annealed with 1875 ng random primers (Invitrogen, Cat#48190-011) for 5 min at 65°C followed by holding step at 4°C. Reverse transcription step was performed using 200 U M-MLV reverse transcriptase (Invitrogen, Cat#28025-013) mixed with 30 U RNAse OUT (Invitrogen, Cat#10777-019), M-MLV reverse transcriptase reaction buffer (Invitrogen, Cat#18057-018) and 1 mM final dNTP mix (Invitrogen, Cat#10297-018). Complementary DNA (cDNA) synthesis was carried out at 37°C for 1h followed by 95°C for 5 min cycle and final hold at 4°C. cDNA was diluted 1:2 or 1:4 in nucleases free water before amplification with LightCycler®480 SYBR Green I Master (Roche, Cat#04-707-516-001) using Light Cycler 480 real-time PCR system (Roche). Absolute abundance of genes relative to B2M housekeeping gene expression was calculated based on a cDNA standard curve using Light Cycler 480 Software 1.7 (Roche). A complete list of primer sequences is provided below.

### Human RT-qPCR primers list

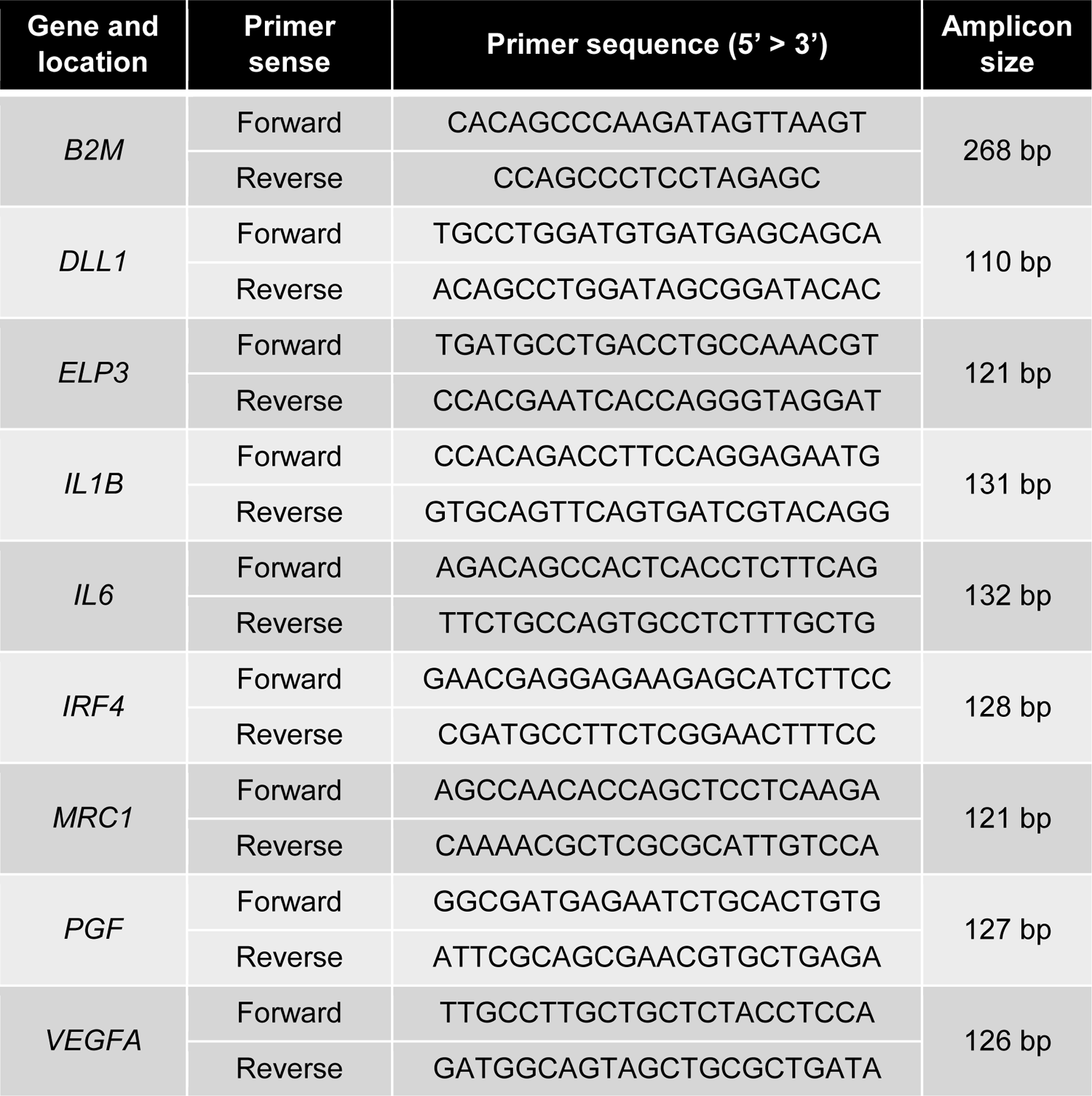

### Mouse RT-qPCR primers list

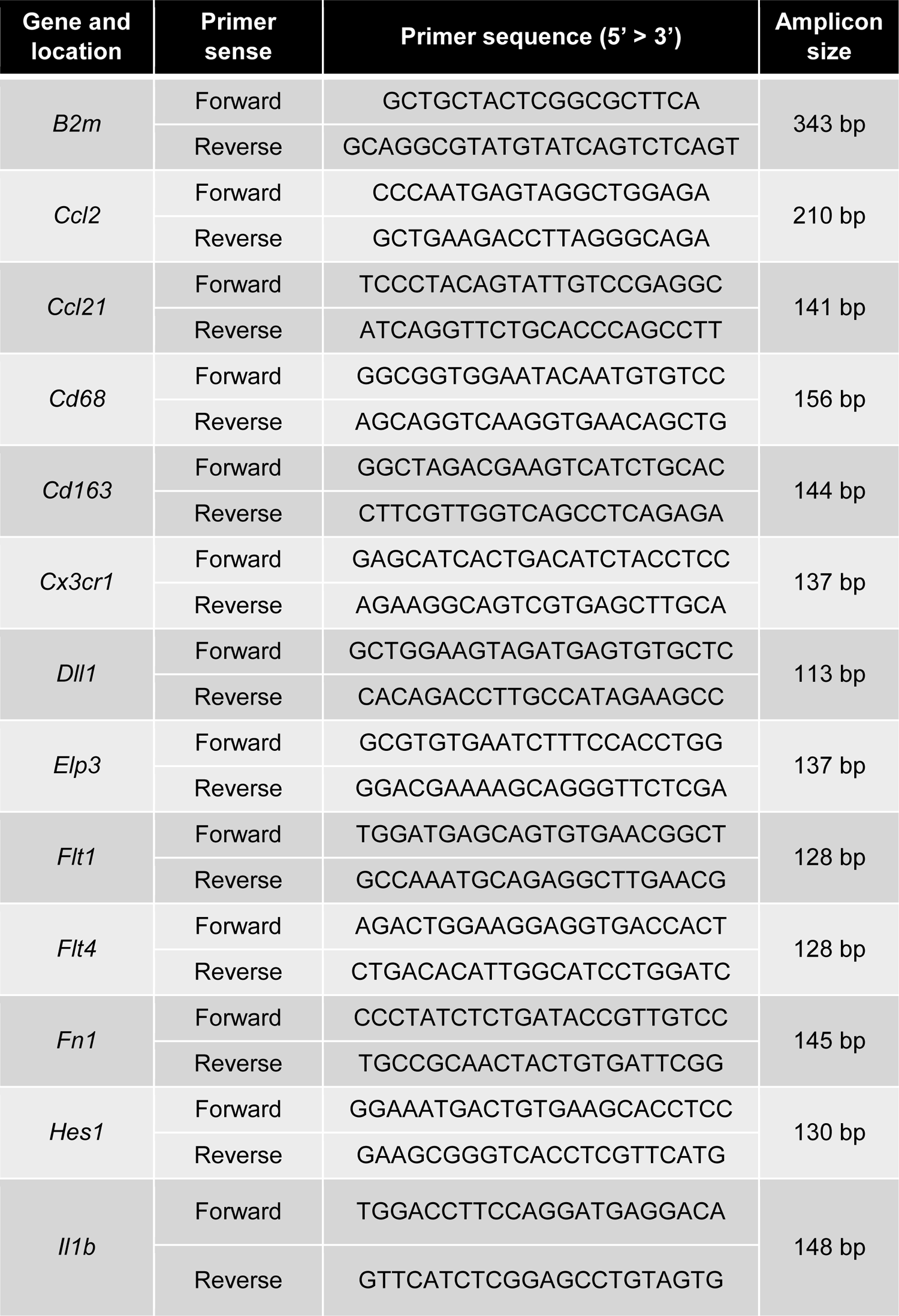
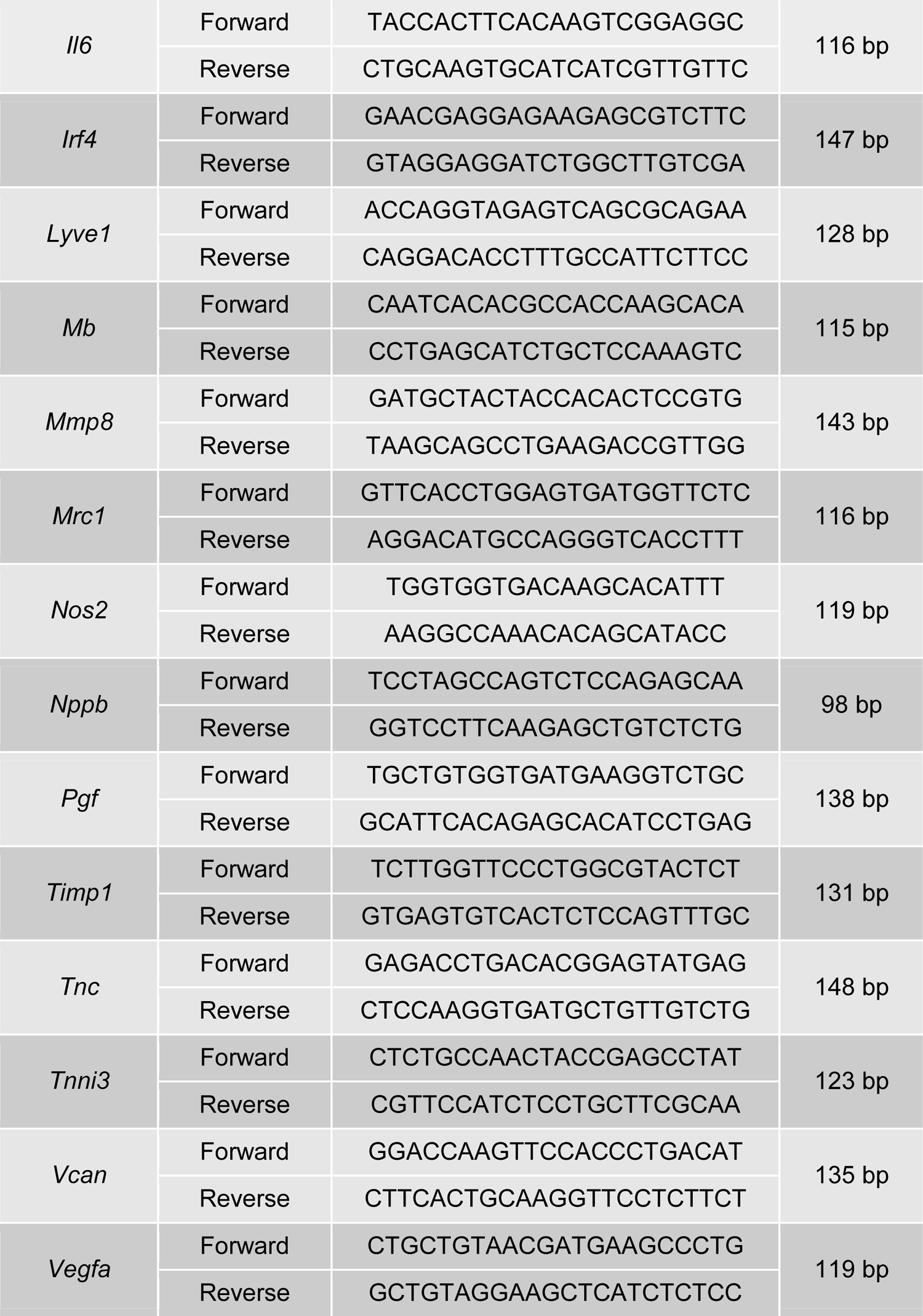

### ChIP-seq data analysis

We downloaded the all sets from the ENCODE portal (Sloan et al. 2016) (https://www.encodeproject.org/) with the following identifiers: ENCSR267NWZ, and ENCSR000ASK respectively for human CD14^+^ monocytes H3K4me3 (GSM1003536) and H3K27me3 (GSM1003564) ChIP-seq data sets.

Reads were aligned to the hg38 genome build using Bowtie2 version 2.3.4.1^45^, ^46^. Unmapped and duplicate reads were removed with Samtools version 1.5^47^. Peak calling was performed with MACS2^48^, ^49^ using default parameters for H3K4me3 samples and with the addition –borad option with H3K27me3 samples. Overlapping peaks were identified with the intersect Bed tool from the bedtools package version 2.26^50^. Annotation of common H3K4me3 and H3K27me3 peaks was performed with PAVIS^51^.

### Native Chromatin Immunoprecipitation (NChIP)

H3K4me3 and H3K27me3 histone modification levels was measured by native ChIP (NChIP) as described^52^. ChIPed DNA was purified by phenol-chloroform extraction, precipitated with ethanol and specific DNA sequences were amplified with LightCycler®480 SYBR Green I Master (Roche, Cat#04-707-516-001) using Light Cycler 480 real-time PCR system (Roche). ChIPed DNA quantity was calculated compared to a genomic DNA standard curve using Light Cycler 480 Software 1.7 (Roche).

### ChIP-qPCR primers list

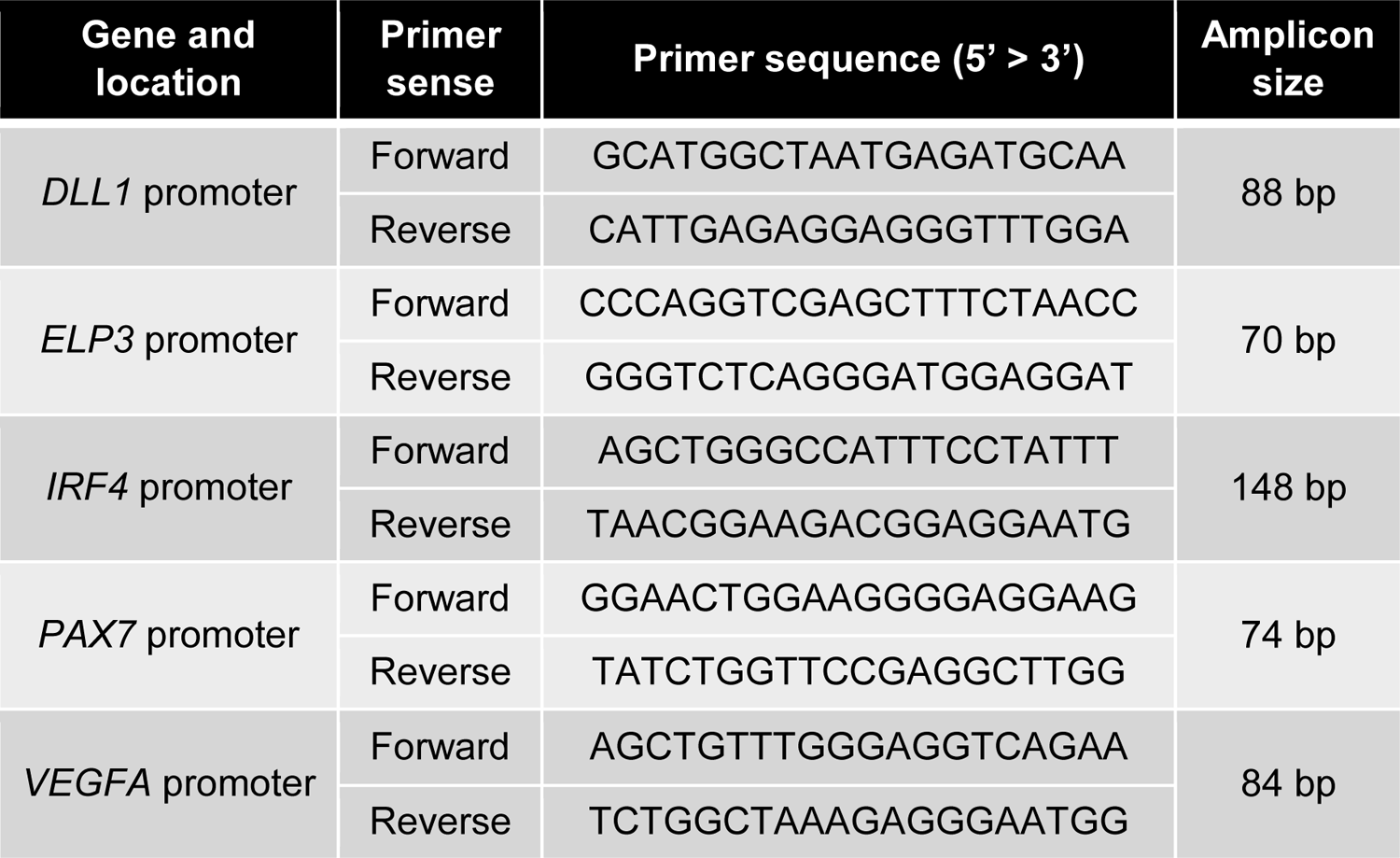

### Heart dissociation

Cardiac samples were collected at 3 or 8 days after MI. Left ventricles (LV) were harvested after perfusion with physiologic serum at 37°C. Infarcted Scar and Border Zones (BZ) were carefully separated and collected in 50ml-falcon tube containing 5mL of RPMI 1640 medium. Three LVs from same group were collected per tube. Digestion enzymes cocktail (collagenase II (5 mg), Dispase (6 mg), DNAse I (300 µg) (Sigma) per 5mL) was added and sample were dissociated through GentleMACS_TM_ (Miltenyi) for 15min. After dissociation, samples were filtered through 70 µm and then 40 µm cell strainer and prepared for FACS staining in PBS-5% FBS buffer.

### Cardiac inflammation analysis by Fluorescence-Activated Cell Sorting (FACS) analysis

Cardiac immune cell infiltration was analyzed by FACS by gating on CD45^+^ leucocytes to exclude endothelial cells and other cardiac cell types from the analysis. On live cells, lymphocytes were gated out by CD3 or CD19 staining. Then, dendritic cells were excluded from the analysis by gating out CD11c^+^ cells. On CD11b^+^CD11c^-^ pool, monocyte/macrophage cell populations were analyzed on their expression of Ly6C versus CX3CR1 staining. Granulocytes were excluded by their lower expression of CX3CR1 compared to monocytes/macrophages subset. Classical/inflammatory and non-classical monocytes/macrophages were defined as Ly6C^hi^/CX3CR1^hi^ and Ly6C^lo^/CX3CR1^hi^, respectively.

### Cardiac functional analysis

Non-invasive echocardiography was performed as described^37^ 3- and 7-days post-MI and GSK-343 treatment. Briefly, mice were anesthetized using Isovet −2% (Osalia, Cat#3248850) and echography was performed with VEVO 3100 ultrasound echograph equipped with a MX550D probe (Fujifilm VisualSonics Cat#51073-45). Short axis view of the left ventricular was performed at the level of papillary muscle and M-mode tracing was recorded. Doppler was used to calculate Velocity Time Integral (VTI) in pulmonary artery. Left ventricular diameter was measured by image analysis using software VEVOLAB V3.1.1.

### Western blotting analysis

TIB-204 nuclei were lysed using Triton Extraction Buffer (TEB: PBS containing 0.5% Triton X 100 (v/v); 2 mM phenylmethylsulfonyl fluoride (PMSF), 0.02% (w/v) NaN3) supplemented with protease inhibitor cocktail before overnight histones acid extraction in 0.2 N HCl at 4°C. Total amount of histones was measured using BioRad protein Assay (Biorad, Cat#500-0006). 2 µg of total histones were resolved by SDS-PAGE electrophoresis followed by transfer on nitrocellulose membrane before hybridation with desired primary antibody previously described overnight at 4°C. Primary antibody signal was amplified using horseradish peroxidase (HRP) coupled specie corresponding secondary antibody.

### Statistical analyses

Data are obtained from at least 3 independent experiments each in duplicate or triplicate for *in vitro* studies. Number of individual animals in each group is indicated for *in vivo* data. Data are expressed as mean values or percentages of control values ± SEM. When indicated, statistical significance between was determined by non-parametric Kruskal-Wallis test followed by Dunns post-hoc test using Graphpad Prism 5 software. The use of non-parametric Kruskal-Wallis test was determined depending Shapiro-Wilk normality test data distribution. Data are considered to be significantly different at values *p* < 0.05. The *p* values are depicted as asterisks (*) or hashtag (#) symbols in the figures as follows: *** or ### *p* <0.001; ** or ## *p* <0.01; * or # *p* <0.05; ns: non-significant.

## Acknowledgments

We thank all physicians from the department of cardiology (Rouen University Hospital) for their help in the collection of peripheral blood samples from patients; Paulus Mulder, Thomas Duflot, Nicolas Perzo and Jérémy Bellien for critical commenting on experiments; Gaëtan Riou from CyFlow flow cytometry and cell analysis facility platform (Institute for Research and Innovation in Biomedicine, IRIB) for his help in performing flow cytometry experiments and analysis; Manon Lefrançois, Florian Vallin and Paul Rouault MSc students for experimental help. We thank the ENCODE Consortium and the Bradley Bernstein, Broad Institute of MIT and Harvard ENCODE production laboratory for generating the GSE29611 ChIP-seq data sets, and more particularly for generating respectively GSM1003536 and GSM1003564 human CD14+monocytes H3K4me3 and H3K27me3 ChIP-seq data sets.

## Author Contributions

JR designed, performed and analyzed all *in vivo* and *in vitro* experiments. DG performed and analyzed echocardiography in mice. SR designed and carried out gene expression experiments. VT performed and analyzed flow cytometry. AD performed and analyzed immunohistochemistry and histology. AC and MB performed ChIP-seq and RNA-seq analysis. JPH carried-out mouse MI model. ZB and CV coordinated collection of peripheral blood samples from cardiology patients. DB and ED designed and coordinated collection of peripheral blood samples from cardiology patients (project EPICAM n°18.07.19.36852 CAT 3, n°-RCB: 2018-A02108-47). DG and VR supervised experiments and provided experimental advice. EB participated in the design of in vivo studies. SF designed, supervised and participated in all *in vivo* and *in vitro* experiments. The article draft was prepared by JR, SF and EB. All authors approved the final version of the article.

