## Supplementary figure S1, S2, S3, S4, S5, S6, supplementary table 4, 5, 6, 7 for "Ezh2 as an epigenetic checkpoint regulator during monocyte differentiation: a potential target to improve cardiac repair after myocardial infarction"

### Supplementary information

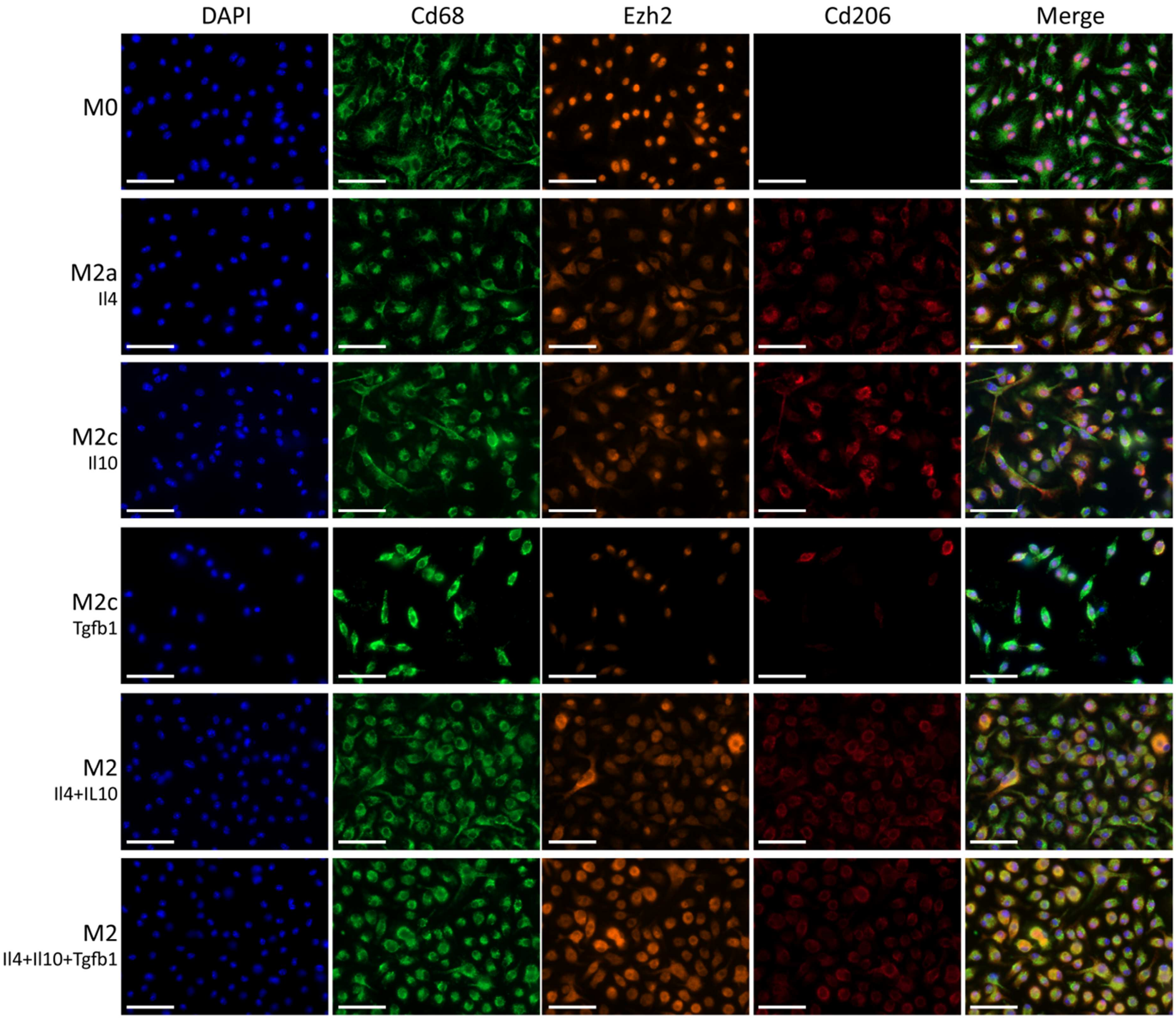


**Supplementary figure S1: Ezh2 cytoplasmic translocation is not restricted to a single M2 macrophage phenotype**

Representative pictures of immunostaining for mature non-polarized M0 macrophages, M2 differentially polarized macrophages with either single Il4 for M2a, Il10 or Tgfb1 for M2c, or combinations of cytokines *in vitro*. Nuclei were stained with DAPI (blue), macrophages with Cd68 (*green*) and M2 polarization was assessed based on Cd206 (*red*) expression. Ezh2 (orange) cellular localization was observed in all cell type but only appeared in M2 macrophages cytoplasm. Scale bars represent 50 µm.

**Supplementary table 1: list of bivalent genes identified by ChIP-Sequencing in CD14+human monocytes**

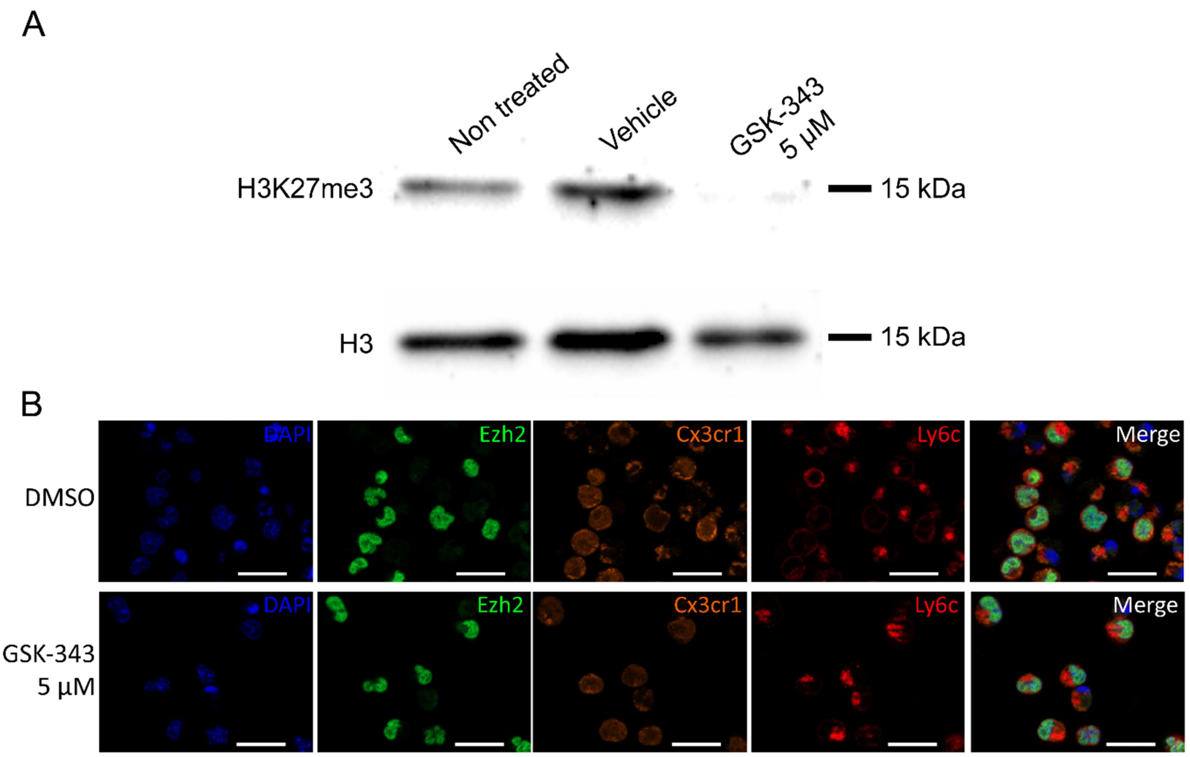


**Supplementary figure S2: GSK-343 reduces H3K27me3 levels without altering Ezh2 expression in murine monocytes**

((A) GSK-343-treatment in murine monocytes (TIB-204 cell line) decreased histone H3K27me3 levels, as determined by western blot. Molecular weights are indicated in kDa. Total histone H3 serves as loading control.

(B) Representative pictures of immunostaining of mouse monocytes treated for 72h with vehicle or GSK-343 (5 µM). Nuclei were stained with DAPI (*blue*), Ezh2 (*green*) cellular localization was analyzed in Cx3cr1 (*orange*) and Ly6c (*red*) expressing monocytes. Scale bars represent 25 µm.

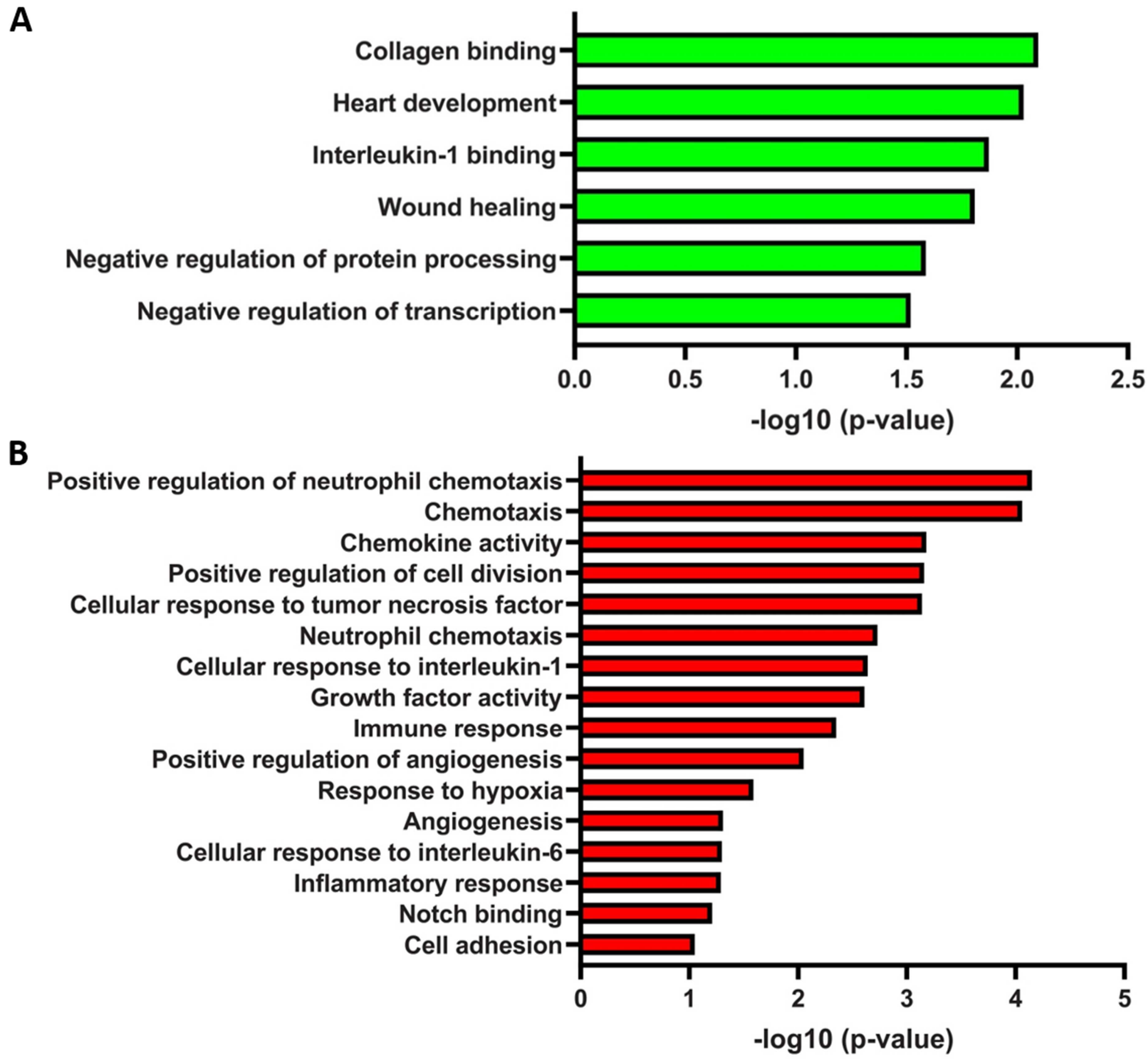


**Supplementary figure S3: GSK-343 treatment of human monocytes promotes expression of genes regulating chemotaxis and angiogenesis**

Representative Gene Ontology (GO) Biological Process categories significantly enriched for down- (A) and up-regulated (B) genes in human monocytes treated with GSK-343 as determined by mRNA-Seq.

**Supplementary table 2: list of down-regulated genes and related GO categories in human monocytes treated with GSK-343**

**Supplementary table 3: list of up-regulated genes and related GO categories in selected circulating human monocytes treated with GSK-343**

| | Group 1<br>Non-CAD<br>n = 20 | Group 2<br>CAD<br>n = 21 | Group 3<br>AMI<br>n = 7 | Overall<br>n = 48 |
|---|---|---|---|---|
| Age (year) | 71±10 | 77±10 | 67±10 | 73±10 |
| Gender (M/F) | 10/10 | 16/5 | 5/2 | 21/17 |
| Erythrocytes ($10^{12}$¢/l) | 4.46±0.50 | 4.50±0.54 | 5.31±0.61 | 4.59±0.61 |
| Leucocytes ($10^{9}$¢/l) | 6.68±1.82 | 7.75±1.79 | 13.22±6.26 | 8.10±3.54 |
| Granulocyte Neutrophils ($10^{9}$¢/l) | 4.22±1.56 | 5.29±1.52 | 10.08±6.03 | 5.55±3.26 |
| Granulocyte Eosinophils ($10^{9}$¢/l) | 0.19±0.15 | 0.20±0.13 | 0.07±0.06 | 0.18±0.14 |
| Granulocyte Basophils ($10^{8}$¢/l) | 0.45±0.24 | 0.51±0.31 | 0.46±0.39 | 0.47±0.29 |
| Lymphocytes ($10^{9}$¢/l) | 1.59±0.70 | 1.62±0.57 | 2.27±1.66 | 1.70±0.86 |
| Monocytes ($10^{9}$¢/l) | 0.63±0.18 | 0.69±0.25 | 0.76±0.30 | 0.68±0.23 |
| Platelets ($10^{9}$¢/l) | 234.9±58.9 | 227.4±51.2 | 279.0±94.1 | 238.1±63.0 |

**Supplementary table 4: patient characteristics**

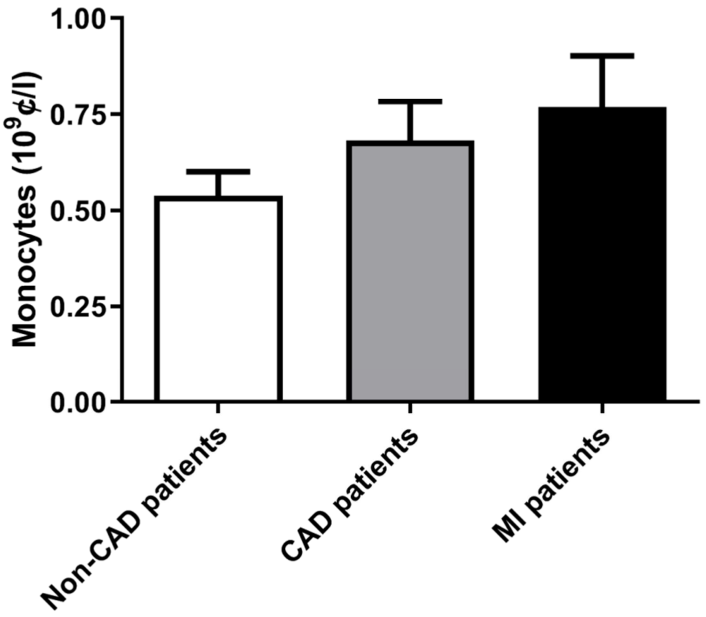


**Supplementary figure S4: Levels of selected circulating monocytes in MI patients as compared to non-CAD and CAD patients**

Blood-derived human monocyte quantification, obtained after negative magnetic selection. Data is represented as mean number of live selected monocytes obtained per ml of collected blood, Non-CAD (n=20), CAD (n=21) and MI (n=7) patients ± SEM.

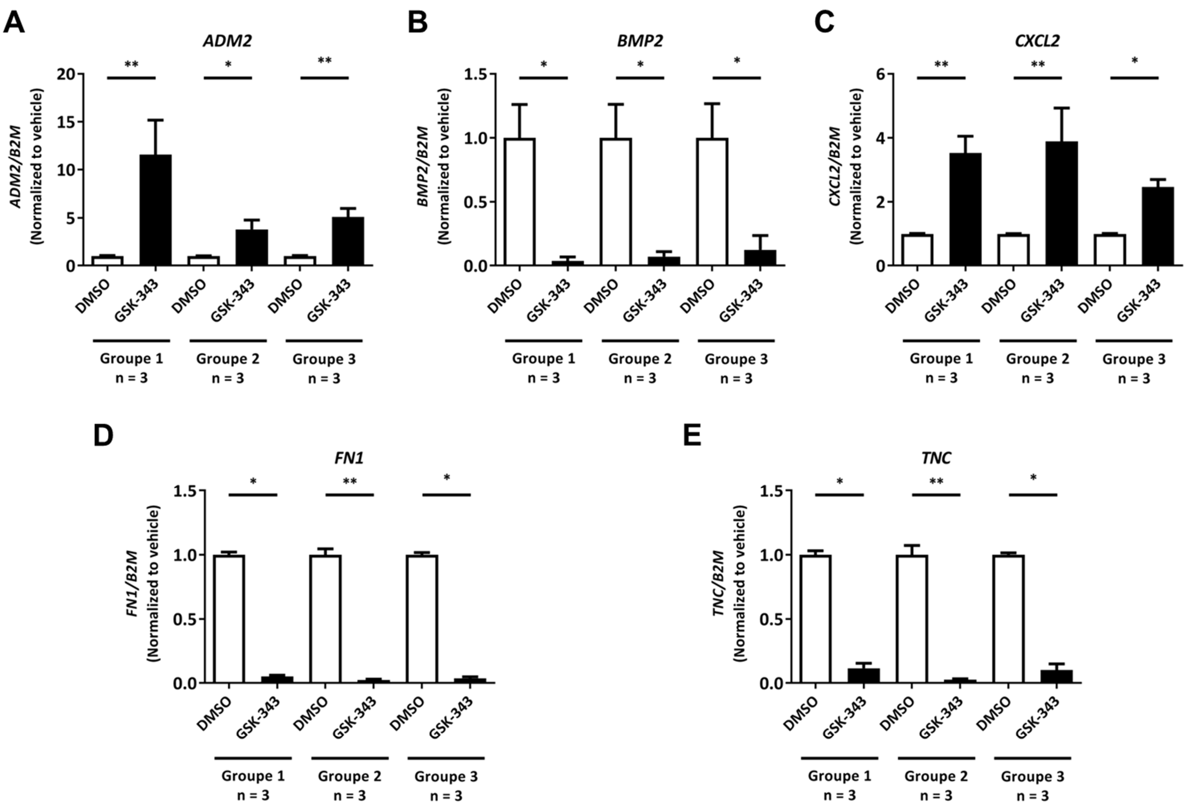


**Supplementary figure S5: Reduced and enhanced expression of genes identified by RNA-seq upon GSK-343 treatment**

Expression of ADM2 (A), BMP2 (B), CXCL2 (C), FN1 (D) and TNC (E) in treated human monocytes from different patient groups with vehicle or GSK-343 was measured par RT-qPCR. Data are presented as mean expression to fold of vehicle treated monocytes ± SEM. Group 1 Non-CAD (n=3), group 2 CAD (n=3) and group 3 AMI (n=3). The p values are depicted as asterisks in the figures as follows: **p <0.01; *p <0.05.

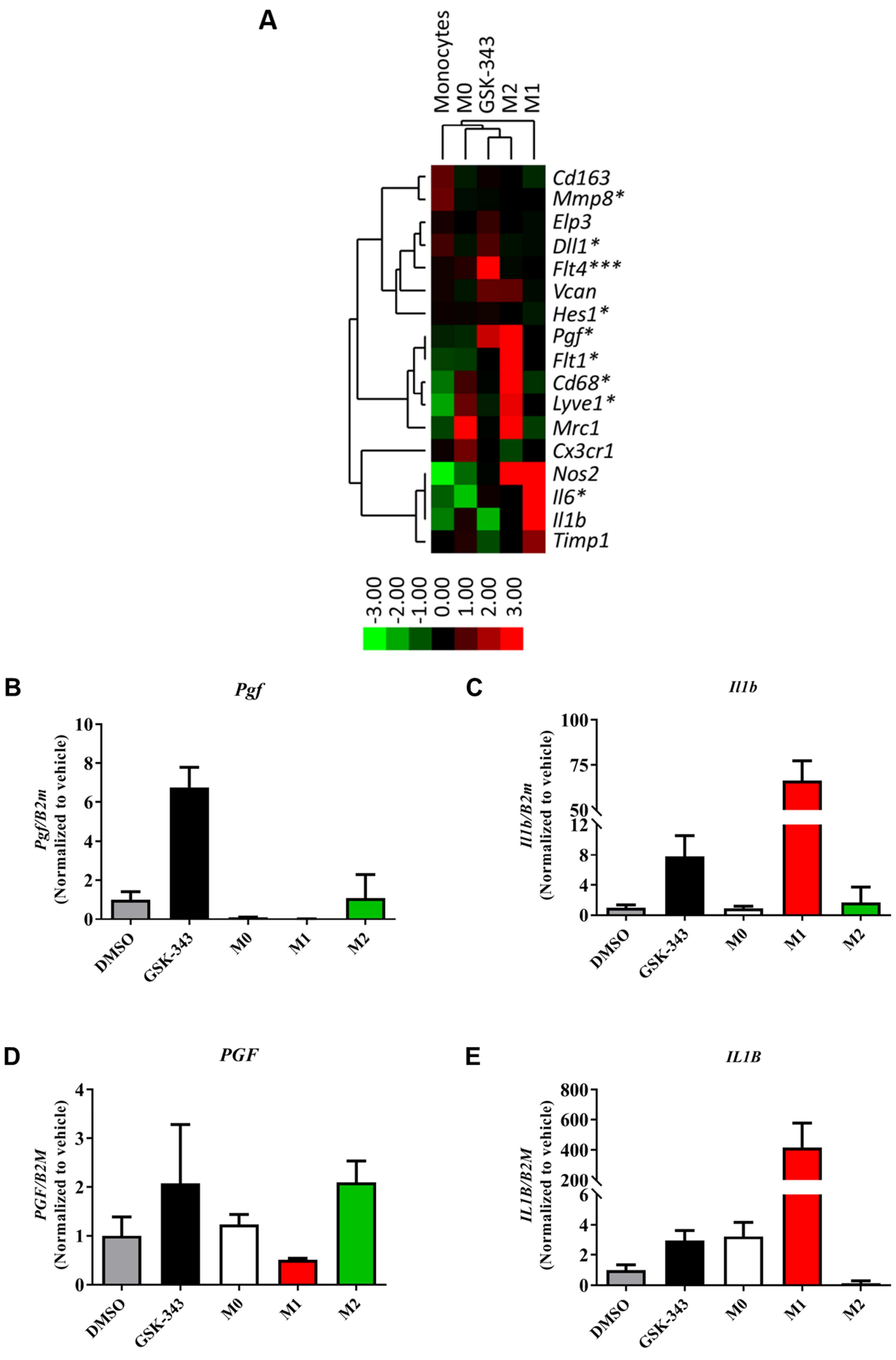
A
Monocytes
M0
GSK-343
M2
M1
Cd163
Mmp8*
Elp3
Dll1*
Flt4***
Vcan
Hes1*
Pgf*
Flt1*
Cd68*
Lyve1*
Mrc1
Cx3cr1
Nos2
Il6*
Il1b
Timp1
-3.00 -2.00 -1.00 0.00 1.00 2.00 3.00
B
Pgf
Pgf/B2m (Normalized to vehicle)
DMSO
GSK-343
M0
M1
M2
C
Il1b
Il1b/B2m (Normalized to vehicle)
D
PGF
PGF/B2M (Normalized to vehicle)
E
IL1B
IL1B/B2M (Normalized to vehicle)

**Supplementary figure S6: EZH2 inhibition with GSK-343 brings monocytes closer to M2 than any other myeloïd cell types**

Gene expression in monocytes treated with vehicle or GSK-343 or following *in vitro* differentiation of monocytes into M0, M1 or M2 macrophages from TIB-204 monocyte mouse cell line (A) or mouse primary circulating selected monocytes (B and C) or human primary circulating selected monocytes (D and E) was measured by RT-qPCR. Data are presented as a targeted transcriptomic array of selected genes for each cell type categories derived from TIB-204 (A) or mean expression reported vehicle treated monocytes ± SEM for mouse primary circulating selected monocytes (B and C) or human primary circulating selected monocytes (D and E). The p values are depicted as asterisks in the figures as follows: ***p <0.001; **p <0.01; *p <0.05; ns: non-significant.

| | 3 days post-MI | | | 8 days post MI | | |
|---|---|---|---|---|---|---|
| | Sham (n=4) | vehicle (n=7) | GSK-343 (n=5) | Sham (n=4) | vehicle (n=5) | GSK-343 (n=8) |
| **Cardiomyocyte sizes (µm²)** | ND | 450.86 ± 28.9 ns | 492.50 ± 12.8 ns | 398.5 ± 19.5 | 546.80 ± 43.80 * | 441.87 ± 23.6 # |
| **Capillary density (Blood vessels/mm²)** | ND | 1681.28 ± 179.3 *** | 1783.40 ± 228.6 ns | 2439.5 ± 179.6 | 1521.0 ± 189.0 *** | 1774.25 ± 199.85 ns |
| **Open lymphatic vessel density (vessels/mm²)** | ND | 7.17 ± 2.09 ns | 5.87 ± 2.02 ns | 5.47 ± 1.25 | 10.34 ± 2.33 p=0.07 | 6.68 ± 1.97 ns |
| **Total macrophages (cells/field)** | ND | 30.58 ± 1.88 * | 38.22 ± 3.26 p=0.11 | 23.48 ± 1.77 | 38.98 ± 4.16 *** | 39.76 ± 5.05 ns |

**Supplementary table 5: Histologic cardiac parameters**

Quantitative analysis of histologic cardiac parameters in border zone (BZ) of sham, vehicle and GSK-343 treated mice at 3- and 8-days post-MI. Data are presented as means ± SEM. Asterisk (*) and hashtag (#) symbols indicates statistically significant difference compared to sham and vehicle condition respectively after Kruskal-Wallis test. # p <0.05, ns non-significant, * p <0.05*** p <0.001. ND indicates Not Determined values.

| Gene function | Gene name | 3 days post-MI | | | 8 days post MI | | |
|---|---|---|---|---|---|---|---|
| | | sham | vehicle | GSK-343 | sham | vehicle | GSK-343 |
| Inflammation | *Ccl2* | 1.00 ± 0.19 | 4.41 ± 0.47 *** | 4.08 ± 0.37 *** | See figure 6b | | |
| | *Ccl21* | 1.00 ± 0.23 | 0.65 ± 0.04 ns | 0.62 ± 0.07 ns | See figure 6b | | |
| | *Il1b* | 1.00 ± 0.19 | 1.66 ± 0.14 ** | 1.51 ± 0.10 ns | See figure 6b | | |
| | *Il6* | 1.00 ± 0.28 | 3.97 ± 0.32 *** | 5.31 ± 0.58 *** | See figure 6b | | |
| | *Il10* | 1.00 ± 0.21 | 1.62 ± 0.14 * | 1.67 ± 0.14 * | 1.00 ± 0.13 | 1.09 ± 0.11 ns | 0.97 ± 0.10 ns |
| | *Irf4* | 1.00 ± 0.25 | 0.46 ± 0.03 *** | 0.51 ± 0.04 *** | 1.00 ± 0.10 | 0.58 ± 0.03 *** | 0.66 ± 0.04 ** |
| | *Tnf* | 1.00 ± 0.27 | 1.50 ± 0.10 ns | 1.47 ± 0.09 ns | 1.00 ± 0.11 | 1.56 ± 0.12 * | 1.52 ± 0.13 ns |
| Angiogenesis | *Dll1* | 1.00 ± 0.30 | 0.59 ± 0.03 ns | 0.61 ± 0.04 ns | See figure 6b | | |
| | *Vegfa* | 1.00 ± 0.23 | 0.65 ± 0.04 ** | 0.74 ± 0.06 ns | 1.00 ± 0.08 | 0.69 ± 0.03 * | 0.71 ± 0.05 * |
| Fibrosis | *Fn1* | 1.00 ± 0.31 | 5.64 ± 0.65 *** | 6.66 ± 0.81 *** | See figure 6b | | |
| | *TnC* | 1.00 ± 0.29 | 10.63 ± 1.91 *** | 10.92 ± 2.15 *** | See figure 6b | | |
| Cardiac dysfunction | *Mb* | 1.00 ± 0.19 | 0.61 ± 0.03 ns | 0.63 ± 0.05 ns | See figure 6b | | |
| | *Nppa* | 1.00 ± 0.29 | 2.04 ± 0.26 ns | 2.01 ± 0.25 ns | 1.00 ± 0.30 | 2.59 ± 0.38 ** | 2.19 ± 0.29 ns |
| | *Nppb* | 1.00 ± 0.18 | 1.77 ± 0.19 ns | 2.18 ± 0.27 * | See figure 6b | | |
| | *Tnni3* | 1.00 ± 0.20 | 0.61 ± 0.03 *** | 0.63 ± 0.06 ** | See figure 6b | | |
| | *Tnnt2* | 1.00 ± 0.28 | 0.96 ± 0.06 ns | 1.07 ± 0.17 ns | 1.00 ± 0.10 | 0.66 ± 0.03 ns | 0.77 ± 0.06 ns |

**Supplementary table 6: Inflammation, angiogenesis, fibrosis and cardiac dysfunction gene expression**

Cardiac transcript levels of indicated genes measured by RT-qPCR from sham (n=8), or vehicle (n=16) or GSK-343-treated (n=18) mice at 3 and 8 days after MI. Values expressed as mean percentages of sham ± SEM with B2m serving as internal control. Asterisk (*) symbols indicates statistically significant difference compared to sham condition. ns non-significant, * $p < 0.05$, *** $p < 0.001$.

| | 3 days post-MI | | | 7 days post MI | | |
|---|---|---|---|---|---|---|
| | sham | vehicle | GSK-343 | sham | vehicle | GSK-343 |
| LV dilatation index | 0.190 ± 0.01 | 0.166 ± 0.06 ns | 0.180 ± 0.04 ns | See figure 6e | | |
| LVED (mm) | 3.75 ± 0.09 | 4.27 ± 0.10 * | 4.21 ± 0.09 ns | See figure 6f | | |
| LVES (mm) | 2.94 ±0.10 | 3.66 ± 0.12 ** | 3.49 ± 0.11 ns | See figure 6g | | |
| LVFS (%) | 21.49 ± 1.73 | 14.50 ± 1.43 * | 15.94 ± 1.43 ns | See figure 6h | | |
| LVEF (%) | 44.09 ± 3.10 | 30.19 ± 2.76 * | 33.39 ± 2.70 ns | See figure 6i | | |
| Cardiac output (ml/min) | 9.45 ± 1.32 | 9.98 ± 0.94 ns | 10.59 ± 0.90 ns | 13.33 ± 1.05 | 11.07 ± 1.31 ns | 12.04 ± 1.29 ns |
| Heart rate (bpm) | 378.7 ± 18.57 | 394.8 ± 11.66 ns | 406.0 ± 7.41 ns | 410.4 ± 12.34 | 425.2 ± 14.22 ns | 429.6 ± 13.24 ns |
| LV mass (mg) | 72.21 ± 1.61 | 85.32 ± 5.51 ns | 94.65 ± 2.88 ns | 74.98 ± 6.51 | 108.10 ± 5.34 * | 100.80 ± 7.38 ns |
| Stroke volume (µl) | 26.54 ± 2.47 | 24.82 ± 2.36 ns | 26.07 ± 2.05 ns | 32.18 ± 1.97 | 26.92 ± 3.18 ns | 27.91 ± 2.54 ns |

**Supplementary table 7: Echocardiographic cardiac dysfunction parameters**

Quantitative analysis of echocardiographic cardiac parameters in sham (n=8), vehicle (n=14) and GSK-343 (n=18) treated mice at 3- and 7-days post-MI. Data are presented as means ± SEM. Asterisk (*) symbols indicates statistically significant difference compared to sham condition. ns non-significant, * p <0.05, *** p <0.001.
